# Alternative Lengthening of Telomeres and CINSARC are interconnected toward non-translocation-related sarcomas progression

**DOI:** 10.64898/2026.01.23.701253

**Authors:** Gaëlle Pérot, Clémence Guerriau, Natacha Roussel, Emeline Sarot, Carine Valle, Pascal Pomiès, Franck Tirode, Delphine Poncet, Frédéric Chibon

## Abstract

Alternative lengthening of telomeres (ALT) is a telomere elongation mechanism activated during oncogenesis and primarily acting in tumors of mesenchymal origin. Although the proteins involved in the machinery enabling ALT to elongate telomeres are becoming better understood, the underlying biology of this mechanism remains unclear. In the present study, we took advantage of a fully characterized cohort of 98 leiomyosarcomas (LMS) from the French Sarcoma Group to further our understanding of the ALT mechanism. We first compared the transcriptomic profiles of ALT^+^ and TERT^+^ LMS and demonstrated a strong enrichment of the CINSARC signature in ALT^+^ tumors. The establishment of an ALT^+^-related signature in these LMS confirmed the close association between CINSARC and ALT in two additional cohorts of non-translocation-related sarcomas. *In vitro* mesenchymal models of spontaneous ALT induction showed increased CINSARC expression following acquisition of the ALT mechanism. Conversely, ALT inactivation, through *BLM* inhibition, led to decreased CINSARC expression. These results establish CINSARC as a new hallmark of the ALT mechanism in non-translocation-related sarcomas and demonstrate the association of a cellular biological process with the CINSARC prognostic signature, namely the ALT mechanism.

## Introduction

Alternative lengthening of telomeres (ALT) is one of two mechanisms, alongside telomerase reactivation, engaged by cancer cells during oncogenesis to maintain telomeres length and survive indefinitely (1,2). First described in yeast, the ALT mechanism is a telomerase-independent pathway that diverts homologous recombination pathways to elongate telomeres (3). ALT telomeres elongation is temporally regulated during the cell cycle and can be detected in late S phase, but primarily in G2 and mitosis (1). ALT is also spatially regulated, occurring in the nucleus mostly at ALT-associated PML bodies (APB), where telomeres cluster in the presence of the recombination machinery (1). In yeast, two ALT pathways have been described: type I, which is RAD51-dependent, and type II, RAD51-independent, both involving homologous recombination processes (4). In humans, two pathways have also been identified: a RAD52-dependent pathway and a RAD52-independent-RAD51-dependent pathway. The exclusivity of these pathways remains controversial, even in yeast, as they can coexist within the same cells (4). Both pathways rely on BLM and POLD3/4 proteins (1,5). BLM is a crucial protagonist in the ALT mechanism, driving the process in association with other proteins of the dissolvasome complex (TOP3α, RMI1 and RMI2). BLM promotes the formation of APBs, remodels the replication fork, participates in the generation of extrachromosomal telomeric repeats (C-circles), dissolves recombination intermediates and recruits endonucleases for 5’ end resection (5).

While 85 to 90% of cancers activate telomerase to maintain telomere length, tumors of mesenchymal origin, particularly non-translocation-related sarcomas, preferentially use the ALT mechanism (1,2). Soft-tissue sarcomas are aggressive (up to 50% of metastases) and heterogeneous tumors (6). They are classified according to the differentiation of the tumor cells with more than 100 different histological subtypes described. From a genetic standpoint, two major groups are defined: on one hand, sarcomas with a simple genetics harboring a recurrent and specific alteration (mutation or translocation), and on the other hand, sarcomas with complex genetics. These tumors are marked by highly rearranged genomes with quasi-systematic p53 and RB1 pathways inactivation, a low mutational burden, frequent whole genome doubling events (7–9) and a high prevalence of the ALT mechanism, ranging from 25% in liposarcomas (10), 47% in osteosarcomas (11), 65% in undifferentiated pleomorphic sarcomas (12) to 78% in leiomyosarcomas (LMS) (8). A recent study showed that the risk of sarcoma is increased by heritable variants of genes involved in telomere biology and mitotic function (13), confirming the importance of these processes in sarcoma oncogenesis. In non-translocation-related soft-tissue sarcomas, the ALT mechanism is associated with chromosomal instability (14) and tumor aggressiveness (14,15), highlighting the need to characterize the telomere maintenance mechanism (TMM) in these tumors. Several hallmarks of ALT have been described, enabling the identification of ALT^+^ tumors. Telomere length can be measured, as ALT^+^ cells typically have long telomeres of highly variable length. The presence of APBs in cells indicates activation of the mechanism, as do extrachromosomal circular telomeric DNA (C-circles) and a high incidence of telomeric sister chromatid exchanges (1,2). While interest in the ALT mechanism is growing, little is known about its inception in cancer cells. Mutations in chromatin-remodeling factors (*ATRX* and *DAXX)*, DNA repair genes (*SMARCAL1* and *SLX4IP*), histone variants (*H3.1* and *H3.3)* and amplification of *TOP3A*, have been identified in ALT^+^ tumors (1,2,16,17). However, collectively these alterations account for less than 50% of ALT^+^ tumors, suggesting that additional mechanisms remain to be elucidated.

Although the steps involved in telomere elongation are becoming increasingly well understood, the biological processes that govern the ALT mechanism, along with the factors that regulate it, remain unclear. In this study, we characterized the TMM in a well-defined cohort of LMS (WGseq and RNAseq data), with the aim of comparing the biological pathways associated with ALT or TERT mechanisms in sarcomas, thereby gaining deeper insight into the ALT mechanism and its underlying biology.

## Materials and Methods

### Tumor samples

**Cohort 1**: the 98 LMS used in this study were collected prospectively by the French Sarcoma Group as part of the ICGC program (International Cancer Genome Consortium) as described in (18). RNAseq for the 64 first LMS are publicly available data, previously used in (18). Those 64 LMS are described with the 34 additional LMS in the Table S1.

**Cohort 2**: the 106 non-translocation-related sarcomas were collected as previously described in (19). RNAseq for Ninety-nine samples of this cohort are publicly available data, previously used in (19) and are described with the additional 7 samples in the Table S2.

**Cohort 3**: the 120 non-translocation-related sarcomas used were obtained from publicly available datasets, part of the TCGA project. RNAseq data were downloaded from the National Cancer Institute GDC Data Portal: https://portal.gdc.cancer.gov/. Those cases are presented in the Table S3.

### Hybrid cell lines

Parental myoblast cell lines (MYOE6 and MYOE7) were established from the same myoblasts culture obtained from a healthy patient as previously described in (20). MYOE6 cell line was generated by lentiviral transduction using pCDH-CMV-MCS-EF1a-Hygro (pCD515B-1, System Biosciences, USA) in which E6 cDNA (HPV16, NC_001526.4:7125-7601) was cloned as recommended by the manufacturer. MYOE7 cell line was generated using pCDH-CMV-MCS-EF1a-RFP-Puro (pCD516B-2, System Biosciences) in which E7 cDNA (HPV16, NC_001526.4:7604-7900) was cloned with the same method. Lentiviral transduction was done as previously described (18). Hybrid cell lines were then obtained by the spontaneous fusion of MYOE6 cell line with MYOE7 one. 150,000 cells of each parental cell line were seeded in co-culture (6-well plates) during 72h and spontaneous hybrid cells were selected by the addition of 100 µg/ml of hygromycin B (Thermo Fisher scientific, France) and 1 µg/ml of puromycin (Thermo Fisher Scientific) in the culture medium. After around ten passages of culture all hybrids entered a massive crisis and only few clones went through it. We thus obtained six hybrid cell lines (HBT1, HBT2, HBT3, HBA1, HBA2 and HBA3).

### Doxycycline inducible shRNA

The *BLM* shRNAs (Sigma-Aldrich, USA) were cloned into a TET-pLKO-neo vector (Addgene, USA, RRID: Addgene_21916) according to the manufacturer’s recommendations and lentiviral transduction was done as previously described (18). Three *BLM* shRNA were tested but only two were kept according to their inhibition efficiency. Sequences of efficient shRNA targeting *BLM*:

#sh2: 5’-CCGGGCCTTTATTCAATACCCATTTCTCGAGAAATGGGTATTGAATAAAGGCTTTTT-3’
#sh3: 5’-CCGGGTGCAGCAGAAGTGGATTAATCTCGAGATTAATCCACTTCTGCTGCACTTTTT-3’.

### Culture conditions

MYOE6, MYOE7 and hybrid cell lines (pre and post-crisis) were cultured in DMEM 4,5 g/L D-Glucose with Glutamax I (Invitrogen, USA) supplemented with 20% fetal bovine serum (FBS, Sigma-Aldrich) and 1% of penicillin-streptomycin (Sigma-Aldrich). Antibiotics were added to the culture medium according to the cell line: 100 µg/ml of hygromycin for MYOE6, 1 µg/ml of puromycin for MYOE7 and both antibiotics for hybrid cell lines.

Cells with *BLM* shRNA were cultured in DMEM 4,5 g/L D-Glucose with Glutamax I (Invitrogen) supplemented with 20% Tetracyclin free fetal bovine serum (TET-FBS, Dutscher, France), 1% of penicillin-streptomycin (Sigma-Aldrich), 100 µg/ml of hygromycin, 1 µg/ml of puromycin, 800 µg/mL of geneticin (Thermo Fisher Scientific). Expression of shRNA was induced by doxycycline (5 μg/mL, Sigma-Aldrich).

All cell lines were cultured in a humidified CO_2_ incubator at 37°C. All cell lines were tested regularly for mycoplasma contamination using the PlasmoTest™ Mycoplasma Detection Kit (Invivogen, France, Cat# rep-ptrk) according to the manufacturer’s recommendations.

### RNA sequencing

Tumors and cell lines RNA extraction, tumors RNA sequencing, alignment and expression quantification were performed as previously described in (18,19).

For cell lines, total RNAseq libraries were prepared with the Stranded Total RNA Prep Ligation with Ribo-Zero Plus Kit (Illumina, USA) using an input of 500ng of Total RNA (DV200 >85.8%). The libraries were profiled with the HS NGS kit for the Fragment Analyzer (Agilent Technologies, USA) and quantified using the KAPA library quantification kit (Roche Diagnostics, France). The dual-indexed libraries were pooled and sequenced on the Illumina NovaSeq6000 instrument (RRID:SCR_016387). More than 50.10^6^ reads per sample were obtained.

For *BLM* shRNA cell lines RNAseq was performed after 10 weeks of doxycycline treatment.

### Bioinformatics analyses

RNA reads were mapped to the hg38 genome assembly with STAR v2.6.0c (RRID:SCR_004463) (21). Low-quality (score < 20) and duplicated PCR paired-end reads were removed with SAMtools v1.8 (RRID:SCR_002105) (22) and PicardTools v2.18.2 (RRID:SCR_006525), respectively. Then, gene expression was quantified with HTSeq v0.9.1 (RRID:SCR_005514) (23).

CINSARC signature score was computed using the ssGSEA method from a previously generated FPKM expression matrix (18). The analysis was applied to each sample with the Revised GSEA R script (https://www.gsea-msigdb.org/gsea) (RRID:SCR_026610), using the 67 genes of CINSARC signature as the sole gene set. For each tumor expression profile, all genes were ranked according to their levels prior to ssGSEA analysis. A normalized enrichment score (NES) was then calculated for each sample using 10 000 unique random permutations of gene order. Tumors were subsequently classified into CINSARC groups as follows: C1 if NES < 0 and p < 0.05 and C2 if NES > 0 and p < 0.05. Samples with p > 0.05 were considered not significant.

Differential gene expression analysis was performed using R (v 4.1.3) package DESeq2 v1.34.0 (RRID:SCR_015687) (24), using the median-of-ratios method for normalization. Gene Ontology analysis (RRID:SCR_002811) was performed using GORILLA (https://cbl-gorilla.cs.technion.ac.il), considering all GO domains. Gene Set Enrichment Analysis was performed using GSEA software v4.2.3 (RRID:SCR_003199) (25). The CINSARC and ALT^+^ signature gene sets were added to the Hallmark gene sets to enable the calculation of adjusted p values (FDR q values). For tumor cohorts, the default recommended parameters were used. For cellular models (hybrid cell lines and BLM inhibition models), gene set permutation was applied, as recommended for small sample sizes.

Figures (Volcano plots, UpSet plots, Dot plot) were done with ggplot2 package v3.5.2 (RRID:SCR_014601), heatmaps were done using pheatmap package v1.0.12 (RRID:SCR_016418).

### TERF2 PML co-immunofluorescence

TERF2 PML co-immunofluorescence results for the sixty-two LMS were previously presented in (18). Cell lines staining was performed using the same antibodies as previously described in (18) and the protocol described in (26). Slides were analyzed using a confocal Zeiss LSM880 Airyscan (RRID:SCR_020925, Zeiss, Germany) and the Zen software (RRID:SCR_013672, Zeiss). To assess the percentage of cells presenting APBs in their nucleus at least ten fields were analyzed totaling a minimum of 100 nuclei counted.

### C-Circle Assay

C-Circle experiment was realized on the 98 LMS. DNA was extracted as previously described in (18). C-circles evaluation and relative quantification of telomeric sequence by TeloPCR were done as previously described (27). A tumor was considered as ALT^+^ by C-circle assay: if CC>4 or if TL>2 and CC≥2 or if TL>1.8 and CC>1.3. A tumor was considered as TERT^+^ by C-circle assay if: TL≤1.6 and CC≤1.3 or if TL≤1.1 and CC≤2. All other cases are considered as ambiguous (Table S1). CC: C-circles, TL: normalized telomere length.

### Tumors TMM status definition

For each tumor the TMM status was defined as followed: all tumors presenting APBs and an ALT^+^ status by C-circle assay were considered as ALT^+^ tumors. All tumors with an ambiguous C-circle result had APBs, so they were considered as ALT^+^ tumors. Tumors with an ALT^+^ result by C-circle assay were considered as ALT^+^ tumors even if no APB were observed on TMA. TERT^+^ tumors were thus those with no APB and a TERT^+^ result by C-circle assay. For tumors with a result obtained only by C-circle assay: only tumors with an ALT^+^ or a TERT^+^ clear result were kept and their TMM was defined according to the result of this experiment (Table S1).

### TRAP assay

Telomerase activity was assessed for each cell line in triplicates using the TRAPeze® RT Telomerase Detection kit (Millipore, USA, Cat# S7710) according to the manufacturer’s recommendations. Real-time RT-PCR amplification was performed using a StepOnePlus^TM^ device (RRID:SCR_015805) and data were analyzed with the StepOne^TM^ software (RRID:SCR_014281, v2.3, Applied Biosystems, USA). Telomerase units were extrapolated from the standard curve of TSR8 control generated by 1:10 serial dilutions (40-0.04 amoles).

### Real-Time Quantitative Reverse Transcription-PCR

Real-time Quantitative RT-PCR was performed as previously described (28), using TaqMan^TM^ Gene Expression assays (Applied Biosystems): Hs00172060_m1 for *BLM*; Hs99999903_m1 for *ACTB* and Hs99999902_m1 for *RPLP0*. Results were calculated as previously described (28).

### Western Blot

Proteins extraction, electrophoresis and secondary antibodies are described in (18). Primary antibodies used are: anti-BLM (Thermo Fisher Scientific Cat# A300-110A, RRID:AB_2064794, 1:1000, 4°C overnight) and anti-beta-actin (Sigma-Aldrich Cat# A5316, RRID:AB_476743, 1:10000, 1h at room temperature). Signal was detected using a Chemidoc Imager (RRID:SCR_019684, Bio-rad, USA) and protein quantification was performed using Image Lab software (RRID:SCR_014210, v6.1, Bio-rad).

### Cell proliferation assay

For hybrid cell lines, 1,200 cells were seeded in six replicates in three 96-well plates. The number of cells was checked by flow cytometry before seeding. At days 3, 4 and 5, cells were washed, trypsinized, and harvested in a final volume of 100 μL of PBS containing 5% of fetal bovine serum (FBS) and 2 mM EDTA (Sigma-Aldrich). The number of viable cells was evaluated by flow cytometry (MACSQuant Analyzer 10, RRID:SCR_020268, Miltenyi Biotec, Germany) based on their morphological features (FSC-A/SSC-A). Data were analyzed using FlowJo (RRID:SCR_008520, v10.10.0, FlowJo LLC, Becton Dickinson, France) and GraphPad Prism (RRID:SCR_002798, v6.01, GraphPad Software Inc., USA) software. Experiment was realized three independent times.

For *BLM* shRNA cell lines, 1,200 cells were seeded in ten replicates in one 96-well plate. The number of cells was checked by flow cytometry before seeding. Medium was changed the next day and doxycycline was added in five of the ten replicates for each cell line. Medium was then changed again at day 3. Cells were harvested and the number of viable cells was evaluated at day 5 post DOX addition as previously described. Experiment was realized three independent times.

### Propidium iodide labelling

Cell lines were resuspended in PBS and fixed with three volumes of absolute ethanol overnight at −20°C. Cells were washed and a solution of propidium iodide (50 μg/mL, Sigma-Aldrich) and RNAse A (25 μg/mL, Thermo Fisher scientific) was added for 15 min at 37°C. DNA content was quantified with propidium iodide fluorescence intensity by flow cytometry (MACSQuant VYB, Miltenyi Biotec) and analyzed using the cycle tool of the FlowJo software (RRID:SCR_008520, v10.10.0, FlowJo LLC) and GraphPad Prism (RRID:SCR_002798, v6.01, GraphPad Software Inc.) software. Experiment was realized in duplicates three independent times.

### Statistical Analyses

Each experiment was repeated three independent times. For examining the statistical significance of the results, analyses were performed with GraphPad Prism (RRID:SCR_002798, v6.01, GraphPad Software Inc.) software. Normal distribution of data sets was examined with a Shapiro-Wilk normality test if relevant. If data passed normality test, statistical significance between two conditions or more conditions was assessed with an unpaired t-test (with Welch’s correction if different SD, two-tailed p-values) or an ANOVA (two-way ANOVA with Sidak’s multiple comparisons test), respectively. Otherwise a Mann-Whitney test (for two groups, two-tailed p-values) was used. Each statistical test used is indicated in each figure legend if relevant with the number of samples and p-values obtained.

## Results

### Transcriptomic analysis reveals CINSARC signature enrichment in ALT^+^ LMS

APB labelling and C-circle assay were applied to characterize the TMM in a cohort of 98 leiomyosarcomas (Table S1), identifying 11 TERT^+^ LMS (11.2%) and 87 ALT^+^ LMS (88.8%). Gene Set Enrichment Analysis (GSEA) revealed that ALT^+^ tumors are strongly associated with cell cycle and DNA repair processes, as evidenced by enrichment of E2F-targets, G2/M checkpoint, mitotic spindle and DNA repair gene sets (Fig. 1A; Table S4). In contrast, no relevant gene set was significantly enriched in TERT^+^ tumors (Table S4).

**Fig. 1:**
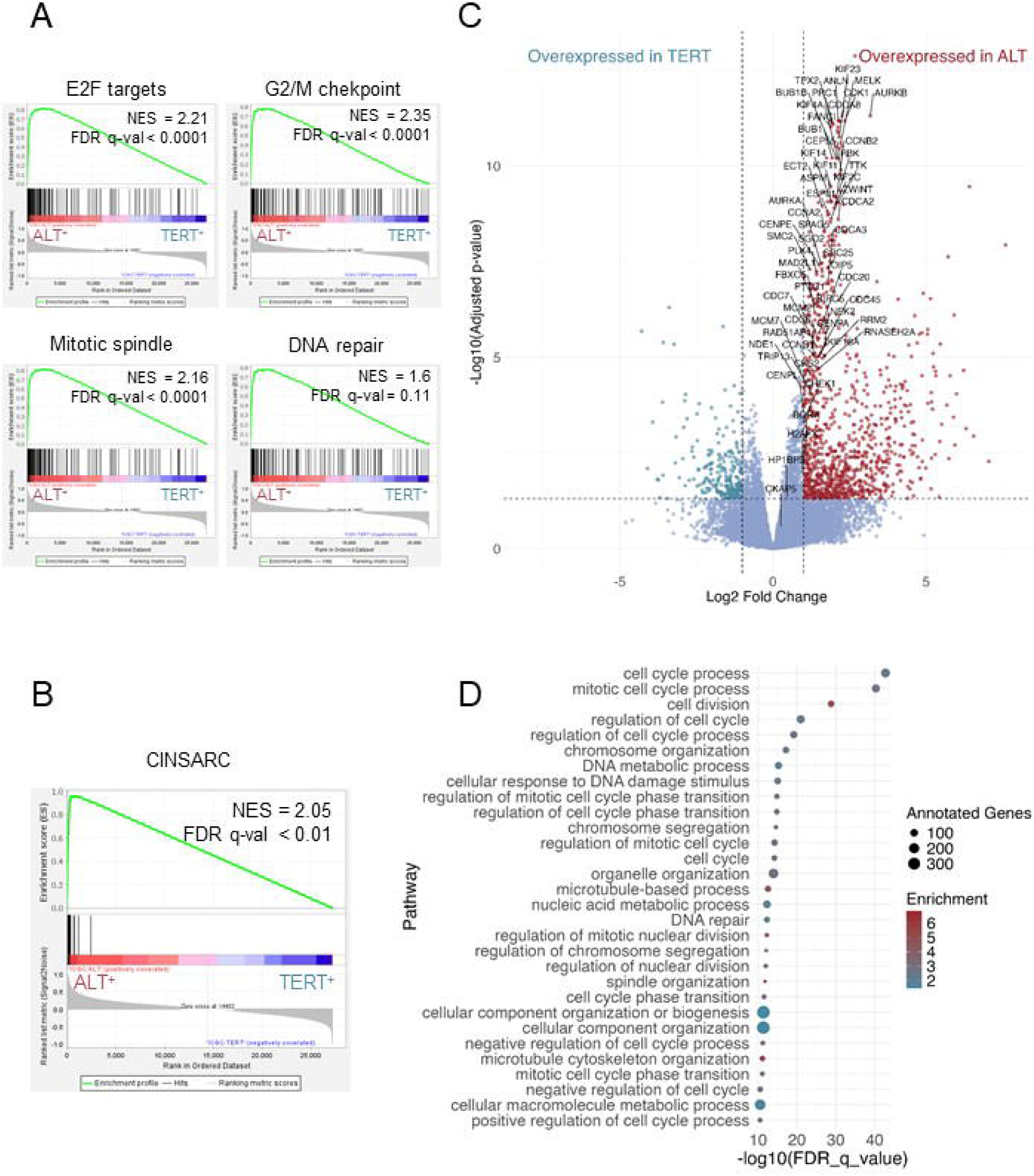
CINSARC gene signature is enriched in ALT^+^ LMS. **A.** Enrichment plots of the 3 most significantly enriched Hallmark gene sets and of DNA repair gene set in ALT^+^ LMS. **B.** Enrichment plot of the 67 CINSARC genes in ICGC LMS (ALT^+^ vs TERT^+^). **C.** Volcano plot showing the differentially expressed genes (DEG) between ALT^+^ and TERT^+^ LMS. Dotted lines highlight thresholds used to define DEG: adj. p-value < 0.05 (Benjamini-Hochberg correction) and absolute log2Fold change >1. Red dots present the 1051 upregulated genes in ALT^+^ LMS and green dots the 239 upregulated genes in TERT^+^ LMS. CINSARC genes are annotated on the figure. **D.** Dot plot representation of Gene Ontology terms enriched for ALT^+^ upregulated genes. Only the 30 most significantly enriched pathways are presented (Table S6). The filled color of each circle indicates the enrichment score. The size of the circle depends on the number of annotated genes enriched in the pathway.

This enrichment of G2/M checkpoint, mitotic spindle and E2F-targets in ALT^+^ LMS prompted us to examine the CINSARC signature in the two LMS groups. The CINSARC signature, a prognostic marker for metastasis initially described in sarcomas (29), comprises 67 genes all related to the control of chromosome integrity and mitosis. The CINSARC signature allows the identification of two tumor groups associated with differences in metastasis-free survival: C1 tumors, which are associated with a good prognosis and lower CINSARC expression, and C2 tumors, which have a worse prognosis and higher CINSARC expression (Fig. S1). As expected, GSEA using CINSARC gene set revealed a strong enrichment in ALT^+^ LMS compared to TERT^+^ ones (Fig. 1B; Table S4).

Differentially expressed genes analysis (DEG) identified 1,051 upregulated and 239 genes downregulated in ALT^+^ LMS (Fig. 1C; Table S5). Among the 1,051 genes upregulated in ALT^+^ LMS, 60 belong to the CINSARC signature (Fig. 1C; Fig. S2 and Table S5). CINSARC genes are overexpressed in ALT^+^ LMS with strong statistical significance (p-value) and moderate fold-change (log2FC), indicating that their expression is consistent and moderately upregulated in the ALT^+^ LMS cohort (Fig. 1C). Forty percent of the CINSARC genes (24/60) are among the top 50 of upregulated genes in ALT^+^ LMS, and 67% (40/60) are in the top 100 (Table S5). Gene ontology analysis of the 1,051 upregulated genes revealed 243 GO terms related to mitosis, replication and DNA repair pathways (Fig. 1D; Table S6). Not only are CINSARC genes associated with these processes, but all genes overexpressed in ALT^+^ LMS are also involved in those pathways related to the S/G2/M phases of the cell cycle. In contrast, no GO terms are enriched among the 239 genes overexpressed in TERT^+^ tumors.

TERT^+^ LMS are highly enriched in CINSARC C1 tumors (10/11 TERT^+^, 91%, C1 corresponding to a low CINSARC enrichment score), while ALT^+^ LMS are predominantly CINSARC C2 tumors (80/86 ALT^+^, 93%, C2 corresponding to a high CINSARC enrichment score) (Table S1), indicating a significant difference in the distribution of CINSARC C1 and C2 tumors between ALT^+^ and TERT^+^ groups (Fisher’s exact test, OR = 117.63, IC95% [13.43; 5595.60], p = 6.56 × 10^−9^). Thus, CINSARC is strongly associated with the TMM of LMS and sustaining these results we could observe that the more CINSARC is enriched, the more C-circles are present in ALT^+^ tumors (Fig. S3).

### Establishment of an ALT^+^-related gene expression signature

We then asked whether the CINSARC signature is also related to TMM in other non-translocation-related sarcoma cohorts. In the other two cohorts, TMM was not characterized, therefore, we used the LMS cohort to define an ALT^+^ expression signature, by first excluding the CINSARC genes from the DEG analysis results and then by selecting the 100 most significantly overexpressed coding genes in ALT^+^ LMS (Table S7). This signature will allow to classify these new cohorts and test whether this classification is associated with CINSARC.

### ALT^+^ signature and CINSARC in LMS

First we confirmed that this ALT^+^ signature is enriched in C2 LMS (GSEA, Fig. S4A) and classified LMS similarly to the expression of the CINSARC gene set (Figs. S4B and S4C).

Gene Ontology analysis of those 100 genes showed that they are involved in the same pathways as the CINSARC genes, *i.e.* mitotic cell cycle process, chromosome segregation and DNA replication (Fig. S5A and Table S8). Moreover, a STRING protein interaction analysis (30) revealed that the majority of the proteins encoded by these 100 genes have predicted and/or validated interactions with proteins encoded by CINSARC genes (Fig. S5B).

Consequently, we then applied this ALT^+^ signature to two additional non-translocation-related sarcoma cohorts to test the hypothesis of an association between CINSARC and TMM.

### Expression of ALT^+^-related gene expression signature enables stratification of non-translocation-related sarcomas based on their CINSARC group

The ALT^+^ signature was thus applied to a first cohort of 106 non-translocation-related sarcomas (cohort 2), for which we first defined a CINSARC enrichment score and established the CINSARC C1 and C2 groups (Table S2). The ALT^+^ signature was strongly enriched in C2 tumors, as observed in the LMS cohort (Fig. 2A; Table S9). Hierarchical clustering using the ALT^+^ signature divided the tumors into two main groups: one composed exclusively of C1 tumors, and the other comprising all C2 tumors along with the three C1 tumors which expressed the most the 100 ALT^+^ genes (Fig. 2B). DEG analysis comparing C2 versus C1 tumors revealed that 84% of the ALT^+^ signature genes were among the 770 genes overexpressed in C2 tumors, while only 2% were among the 964 genes overexpressed in C1 tumors (14 genes were not differentially expressed in either group) (Table S10).

**Fig. 2:**
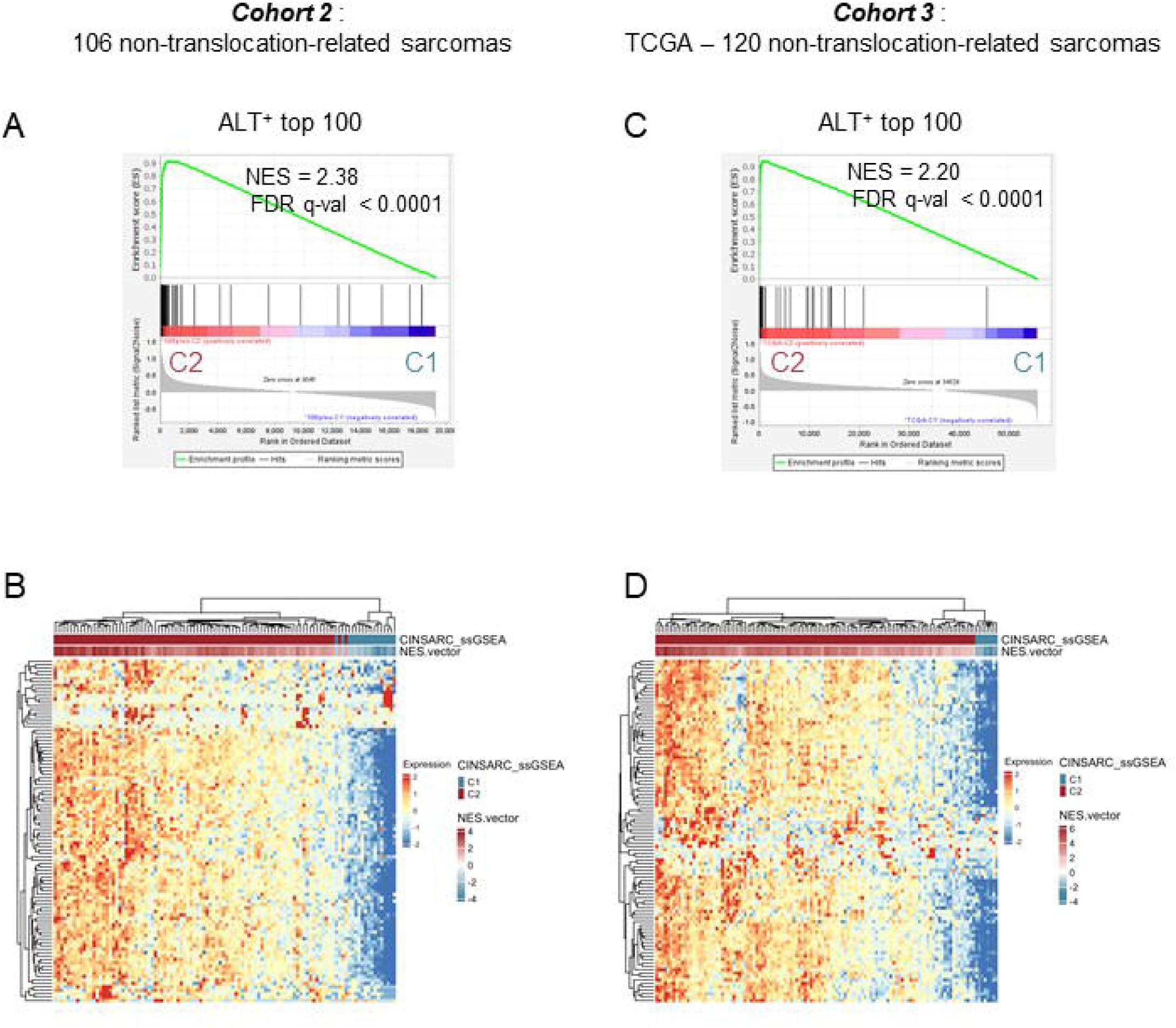
ALT^+^-related gene signature classifies non-translocation-related sarcomas into CINSARC groups. **A.** Enrichment plot of the ALT^+^-related gene signature in the 106 non-translocation-related sarcomas from cohort 2. C1: tumors with a low CINSARC enrichment score and C2: tumors with a high CINSARC enrichment score. **B.** Unsupervised hierarchical clustering heatmap of tumors of cohort 2 using the ALT^+^-related signature. CINSARC ssGSEA groups (C1 and C2) and CINSARC Normalized Enrichment Score (NES) are indicated. **C.** Enrichment plot of the ALT^+^-related gene signature in the 120 nontranslocation-related sarcomas from the TCGA cohort (C2 versus C1 tumors). **D.** Unsupervised hierarchical clustering heatmap of tumors of TCGA cohort using the ALT^+^-related signature. CINSARC ssGSEA groups and NES are indicated.

We then proceeded in the same way for 120 non-translocation-related sarcomas from the TCGA sarcoma cohort (Table S3). We observed that the ALT^+^ signature was also strongly enriched in C2 tumors in this third cohort (Fig. 2C; Table S11). Hierarchical clustering using the ALT^+^ signature split the tumors into two groups: one with high expression of the signature, composed exclusively of C2 tumors, and another including C2 tumors with the lowest NES score along with all C1 tumors (Fig. 2D). DEG analysis revealed that in this third cohort, 92% of the ALT^+^-signature genes were among the 1183 genes overexpressed in C2 tumors, while none were among the 506 C1 overexpressed genes (Table S12). Overall, 80% of ALT^+^-related genes were found to be overexpressed in C2 tumors across the two additional cohorts.

Analyzing more broadly the overexpressed genes across the three cohorts, we observed that only ALT^+^_ICGC_/C2_cohort2&TCGA_ tumors shared a set of commonly overexpressed genes (252 genes, including 59 and 80 genes of the CINSARC and ALT^+^ signatures respectively) (Fig. 3A). In contrast, for ALT^+^ /C1, TERT^+^_ICGC_/C1_cohort2&TCGA_ and TERT^+^_ICGC_/C2_cohort2&TCGA_ tumors, there were no genes - or at most one gene - commonly overexpressed across the three cohorts (Figs. 3B-3D).

**Fig. 3:**
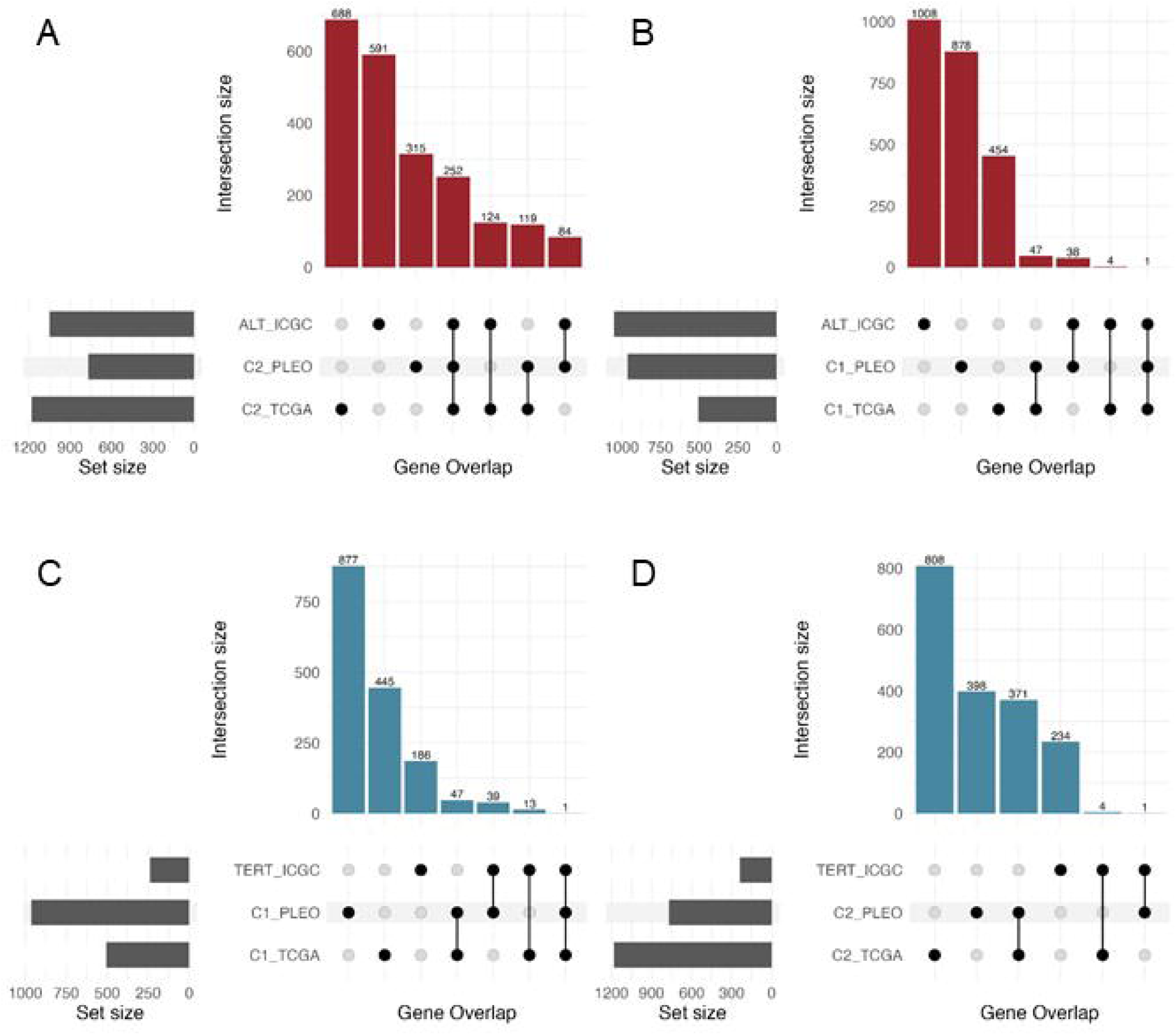
ALT^+^ and C2 tumors shared the higher number of differentially upregulated genes. UpSet plots showing the numbers and interactions of the differentially expressed genes (DEG) between the 3 cohorts (ICGC: 98 LMS, Pleo: cohort 2 of 106 non-translocation-related sarcomas, TCGA: 120 non-translocation-related sarcomas). DEG analysis in ICGC cohort was performed between ALT^+^ and TERT^+^ LMS and in the two other cohorts between C2 and C1 tumors (high and low CINSARC enrichment score respectively). The horizontal bars show the number of DEG in each set, while the vertical bars represent the intersection size. Dark spheres indicate which sets are represented in the vertical bar. **A.** Upset plot between ALT^+^_ICGC_ and C2_cohort2&TCGA_ tumors. **B.** Upset plot between ALT^+^_ICGC_ and C1_cohort2&TCGA_ tumors. C. Upset plot between TERT^+^_ICGC_ and C1_cohort2&TCGA_. **D.** Upset plot between TERT^+^_ICGC_ and C2_cohort2&TCGA_ tumors.

Interestingly, the CINSARC genes and the ALT^+^-related genes show similar behavior across all studied cohorts: they are consistently overexpressed in C2 tumors with high statistical significance (low p-value) and moderate log2FC, regardless of the cohort analyzed (Fig. S6). The genes that deviate from the typical CINSARC behavior, specifically, those showing a high log2 fold change but a low statistical significance (Figs. S6C and S6D), are cohort-specific. These 20 genes are not consistently overexpressed across all cohorts and contribute, little or not at all, to the CINSARC/ALT^+^ gene network (Fig. S5B).

Altogether, these results suggest a biological link between CINSARC and ALT, supporting the need to functionally investigate this interconnection in cellular models.

### Spontaneous acquisition of the ALT mechanism leads to CINSARC enrichment in cellular models

Non-translocation-related sarcomas oncogenesis is primarily driven by systematic inactivation of the p53 and RB1 pathways, together with a whole-genome doubling event (WGD) in most of cases, ultimately leading to a highly rearranged genome (7). We previously showed that WGD could occur in non-translocation-related sarcoma cell lines trough spontaneous fusion (31). Moreover, we demonstrated that, in a mesenchymal context, cell fusion not only induces genome doubling but also promotes genomic instability and tumor initiation, generating hybrid tumors whose genomes closely resemble those of human non-translocation-related sarcomas (32,33). We thus generated tetraploid sarcoma cellular models (Fig. S7A) through the spontaneous fusion of myoblasts with altered p53 and RB1 pathways (one inactivated pathway per cell; Fig. 4A). After bypassing a replicative senescence crisis, the resulting cells spontaneously activated one of the two TMM mechanisms: three derived cell lines were TERT^+^ (exhibiting telomerase activity and lacking APBs) and three were ALT^+^ (lacking telomerase activity and displaying APBs) (Figs. S7B and S7C). This model therefore enabled the investigation of both TMMs within the same mesenchymal genetic context.

**Fig. 4:**
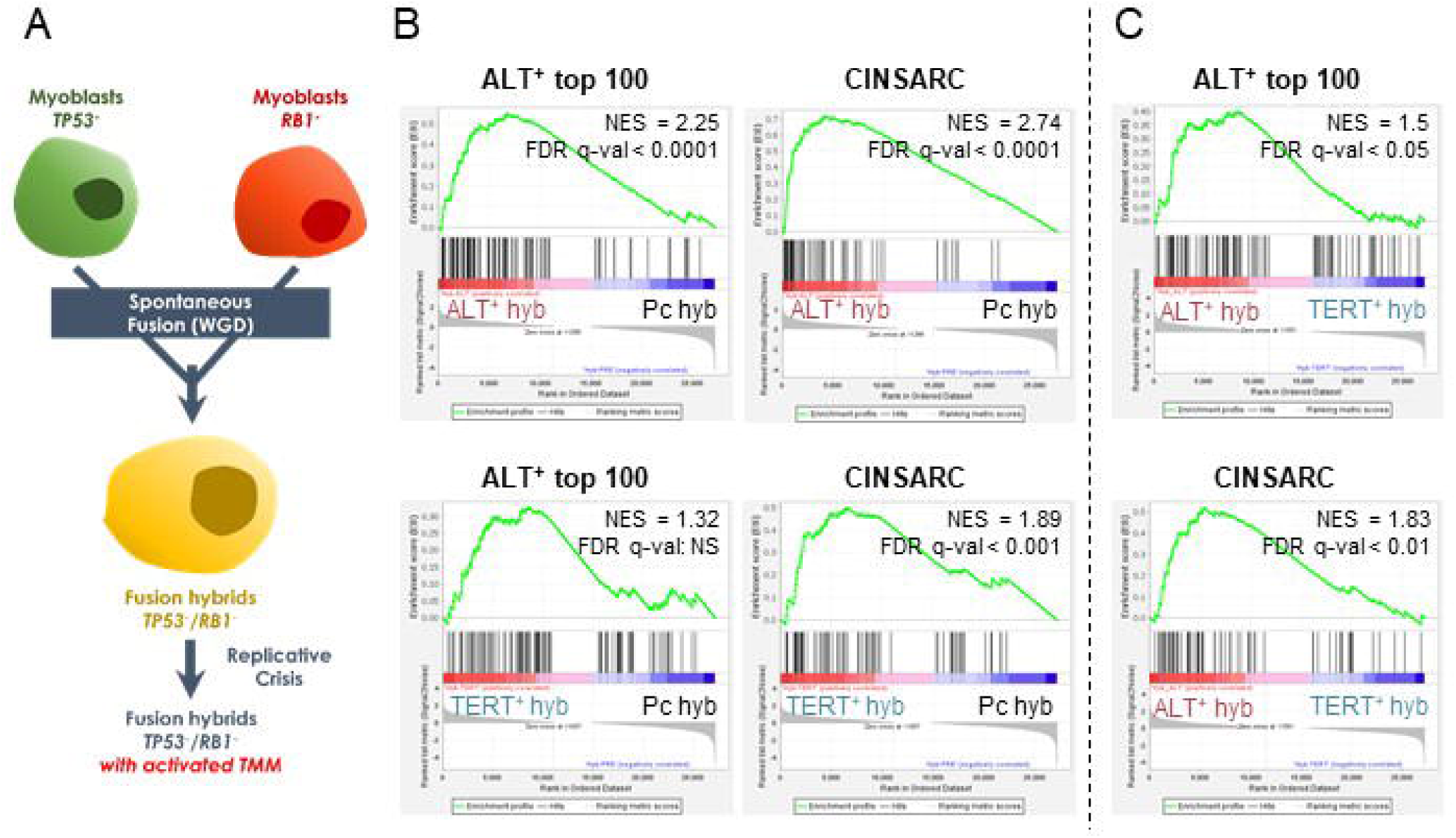
CINSARC signature expression is increased during ALT mechanism acquisition. **A.** Schematic representation of cellular models establishment. WGD: whole genome doubling, TMM: telomeres maintenance mechanism. **B.** Enrichment plots for the ALT^+^-related and CINSARC signatures in ALT^+^ or TERT^+^ hybrids when compared to the hybrids before they activated the TMM (pre-crisis hybrids). Hyb: hybrid cell lines, Pc: pre-crisis, NS: not significant. **C.** Enrichment plots for the ALT^+^-related and CINSARC signatures in ALT^+^ hybrids when compared to TERT ones.

Comparison of hybrid transcriptomes before and after the senescence crisis revealed that ALT^+^ hybrids are enriched for both the ALT^+^ and CINSARC signatures (Fig. 4B; Table S13). While CINSARC is also upregulated in TERT^+^ hybrids, direct comparison shows that enrichment is stronger in ALT^+^ hybrids, consistent with tumors (Fig. 4C; Table S13). This suggests that CINSARC upregulation in ALT^+^ cells is linked to ALT acquisition, whereas in TERT^+^ hybrids it likely reflects increased proliferation.

### ALT phenotype inhibition reduces CINSARC genes expression

*BLM* is a major actor of ALT and several studies have shown that *BLM* inhibition leads to ALT pathway suppression (5,34). *BLM* is overexpressed in ALT^+^ tumors (LMS-ICGC) (Table S5) as well as in ALT^+^ hybrids (Fig. S7D). *BLM* belongs to the ALT^+^ signature and to the 252 genes commonly overexpressed in ALT^+^ and C2 tumors (Tables S5, S7, S10 and S12). Therefore, *BLM* represents a promising target to inhibit the ALT mechanism and to test whether this inhibition reduces CINSARC expression.

Upon *BLM* inhibition by inducible shRNA in the 3 ALT^+^ hybrids (validated at both RNA and protein levels, Figs. S8A and S8B), we observed a strong and significant decrease in the number of APB^+^ cells in all ALT^+^ cell lines compared to controls, which persisted over time (Fig. 5A; Fig. S9). GSEA analysis revealed that gene sets related to G2/M checkpoint, E2F targets, DNA repair and mitotic spindle were enriched in control cells compared to those under *BLM* inhibition, as observed in ALT^+^ LMS (Fig. 5B; Table S14). Similarly, both the ALT^+^ and CINSARC signatures were decreased following *BLM* inhibition (Fig. 5B; Table S14).

**Fig. 5:**
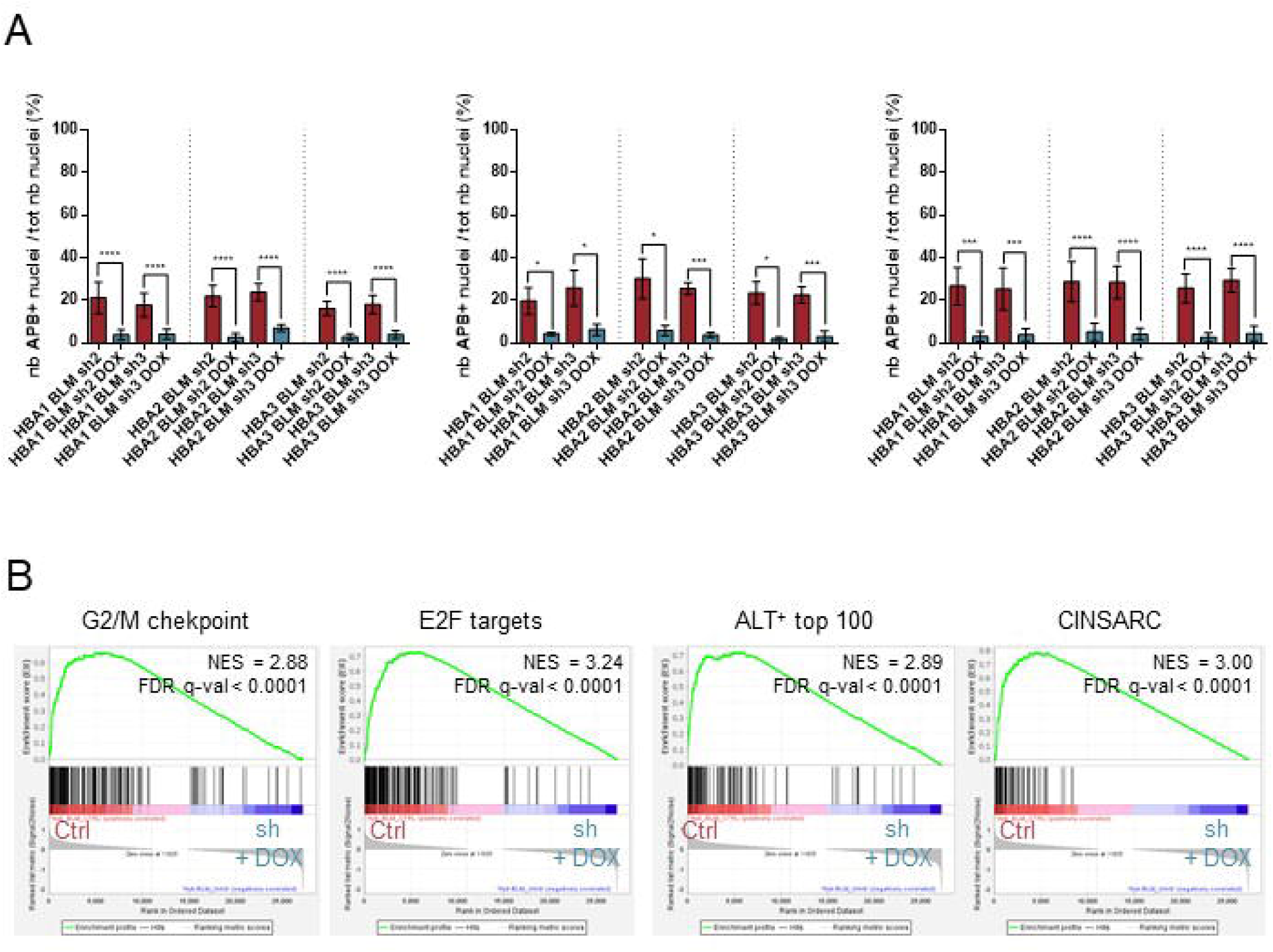
CINSARC signature expression is decreased upon ALT inhibition. **A.** The bar charts present the quantification of the % of cells presenting APBs for each cell line after 72h (left), 4 weeks (middle) and 10 weeks (right) of DOX treatment. Results are mean ± SD. *BLM* inhibition induced by DOX treatment was assessed in parallel by qRT-PCR (72h), qRT-PCR/western blot (4 weeks, Figs. SBA and SBB) and RNAseq (1O weeks). Unpaired t-test with Welch’s correction. *: p<0.05, ***: p<0.001 and ****: p<0.0001. **B.** Enrichment plots for 2 highly enriched Hallmark gene sets, for the ALT^+^-related and CINSARC signatures in ALT^+^ hybrids without *BLM* inhibition (Ctrl, shRNA without DOX induction) compared to the same hybrids with *BLM* inhibition (sh+DOX, shRNA induction by DOX treatment).

Consequently, inhibition of the ALT mechanism leads to a reduction in CINSARC expression, confirming their biological relationship.

### CINSARC expression does not merely reflect increased proliferation

Genes from both signatures are known to have a cell cycle-dependent expression. Interestingly, in the hybrid models, TERT^+^ cell lines proliferate faster than ALT^+^ ones (Fig. 6A; Fig. S10), ruling out the hypothesis that the enrichment of CINSARC and ALT^+^ signatures in ALT^+^ cells is due to or induces higher proliferation.

**Fig. 6:**
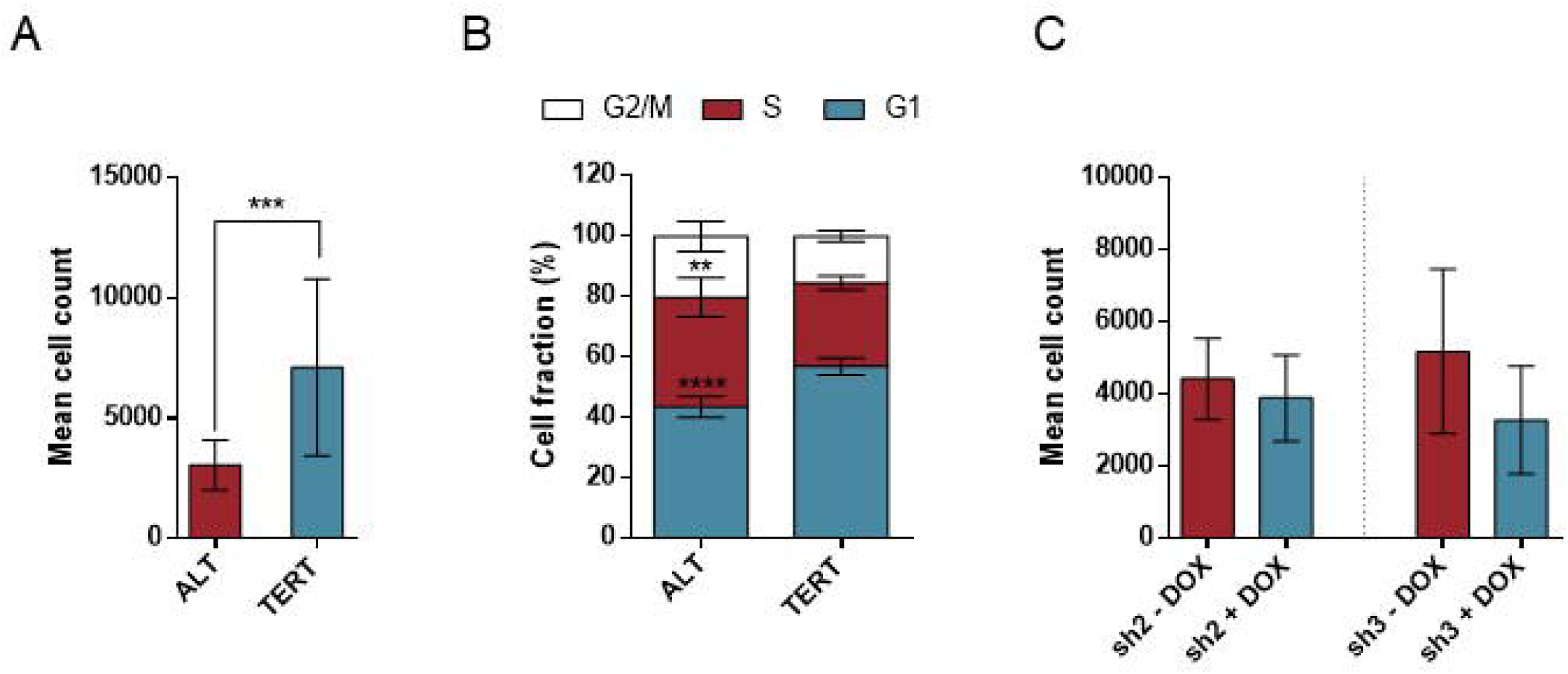
Proliferation assessment in cellular models. **A.** The bar chart corresponds to the mean of three independent proliferation experiments for the 3 TERT^+^ cell lines on one hand and for the 3 ALT^+^ cell lines on the other hand. For each cell line 1200 cells were seeded and proliferation was assessed at days 3, 4 and 5 (Fig. S10). Results presented here show the mean cell count on day 5 ± SD. Mann-Whitney test. ***: p < 0.001. **B.** Stacked bar chart presenting the result of three independent experiments in which cell fractions in G1, S and G2/M phases of the cell cycle were assessed using propidium iodide labelling for 2 TERT^+^ hybrid cell lines and 3 ALT^+^ cell lines. HBT3 TERT^+^ cell line could not be analyzed because of the presence of two populations with different ploidies. Results correspond to the mean ± SD. Two-way ANOVA with Sidak’s correction for multiple comparisons test. **: p < 0.01 and ****: p < 0.0001. Individual data for each cell line are presented in Fig. S11. **C.** The bar chart corresponds to the mean of three independent proliferation experiments for the 3 control cell lines (-DOX, in red) and for the 3 cell lines under DOX treatment (+DOX, in blue). Two shRNAs were used (sh2 & sh3). Results are mean ± SD. Unpaired t-test with Welch’s correction. No significant result. *BLM* inhibition induced by DOX treatment was assessed in parallel by real-time quantitative RT-PCR and western blot (Figs. S12A and S12B). Individual data for each cell line are presented in Fig. S12C.

We observed that ALT^+^ cell lines had significantly fewer cells in G1 and more cells in S/G2/M phases compared to TERT^+^ ones (Fig. 6B; Fig. S11). These higher proportions of cells in S/G2/M phases in ALT^+^ cell lines, despite their lower proliferation rates, likely indicate that the duration of S/G2/M is longer in ALT^+^ cells, which could explain the increased CINSARC expression.

No significant difference in proliferation was observed between control and *BLM*-inhibited cell lines (Fig. 6C; Fig. S12), ruling out the possibility that the reduction in CINSARC genes expression under *BLM* inhibition is due to decreased proliferation. Altogether, these data demonstrate a strong association between the CINSARC signature and the ALT mechanism.

## Discussion

TMM activation represents a critical bottleneck in tumoral transformation, predominantly achieved through ALT in non-translocation-related sarcomas. The data presented here indicate that ALT and CINSARC are interconnected mechanisms and pathways. The key issue arising from this observation is whether CINSARC is part of a process that occurs alongside telomere elongation by ALT, or if CINSARC is ALT itself. This distinction is crucial as CINSARC is now a well-established and validated prognostic marker in sarcomas, yet its underlying biology and relation to tumor aggressiveness remain unexplained.

Most of the CINSARC and ALT^+^-related genes, that are upregulated in ALT^+^/C2 tumors, are connected by a shared biological function (Fig. S13A), suggesting that they all contribute to a common function in non-translocation-related sarcomas biology. In line with these results, previously published data in pediatric osteosarcomas, showed that pathways involving cell cycle checkpoint, chromosome maintenance, DNA replication, G2/M checkpoint, M-phase, pathways that include CINSARC genes, are enriched in ALT^+^ tumors (17). All CINSARC genes are associated with processes occurring during S/G2/M phases of the cell cycle, when telomeres are elongated, raising the question of whether they are directly involved in the ALT process. Notably, four proteins encoded by CINSARC genes have already been identified as participating directly in different stages of the ALT mechanism. These include CHEK1, which regulates BLM recruitment to telomeres during the early steps of ALT; TOP2a, which modulates BLM helicase activity; HB1BP3, which controls chromatin compaction at telomeres during ALT; and RAD51AP1, which promotes, in cooperation with TERRA, the formation of telomeric R-loops and D-loops, homologous recombination intermediates involved in ALT mediated telomere elongation (35–38). Some other CINSARC genes have been described as involved in telomeres biology (*AURKB, BIRC5, CDK1, CDCA8, CDC20, CDCA2, BUB1, BUB1B, TPX2, FOXM1, ESPL1, CENPA, NEK2, RRM2, CDC6, MAD2L1, FBXO5, TRIP13*) or in DNA repair processes, such as Break Induced Replication or DSB repair in response to replication stress, both playing a role in ALT mechanism (*ASPM, AURKA, AURKB, BIRC5, BORA, CCNA2, CDK1, CDC45, CDC6, CDC7, MCM2, MCM7, H2AFX, KIF4A, PBK, PTTG1, TRIP13*) (for references see Table S15).

Not only there is a strong association between ALT and CINSARC expression observed in tumors, but our results from cellular models also demonstrate that CINSARC expression is regulated alongside ALT induction or inhibition. We observed that CINSARC expression increases more in ALT than in TERT hybrids during spontaneous TMM acquisition, even though the latter proliferate faster. While the CINSARC genes could be considered as proliferation markers in TERT hybrids, their role in proliferation is likely not the primary factor in ALT hybrids, since CINSARC expression rises more despite lower proliferation. Supporting this, Jemaà *et al*. showed that sarcoma cell line subclones with higher expression of CINSARC mitotic genes did not exhibit a proliferative advantage compared to those with lower expression (39). Moreover, upon ALT inhibition, CINSARC expression decreases without affecting cell proliferation, suggesting that CINSARC genes role in ALT is proliferation independent. Together, literature data and these functional results suggest that CINSARC genes act directly in ALT. Future studies should aim to determine how each CINSARC gene contributes to telomere elongation or whether they are part of a cellular context that facilitates this process.

All CINSARC genes are described as being temporally regulated and mainly expressed during S/G2/M phases of the cell cycle. In hybrid models, ALT^+^ cells have a higher proportion of cells in S/G2/M phases compared to TERT^+^ cells, despite slower proliferation. Therefore, we hypothesize that the duration of these phases is extended in ALT^+^ cells. The prolonged duration of these phases, when CINSARC genes are expressed, could likely explain the higher expression of the signature in ALT^+^ cells. This suggests that during ALT establishment, the phases associated with telomere elongation may lengthen, facilitating or enabling the mechanism. Alternatively, ALT may be established only in cells that have longer S/G2/M phases. The extended duration of these phases could provide a selective advantage by maintaining the expression of replication, repair and mitosis genes over time, all of which also contributing to the ALT mechanism. Our findings thus open new perspectives to deepen our knowledge of the molecular basis of ALT. Not only is CINSARC a new hallmark of the ALT mechanism, but in soft tissue non-translocation-related sarcomas at least, CINSARC essentially *is* ALT.

This study reveals a strong and coherent biology underlying ALT^+^ tumors. In contrast, TERT^+^ non-translocation-related sarcomas do not appear to share such a unifying program: analysis of genes overexpressed in TERT^+^ LMS or C1 sarcomas fails to highlight any consistently enriched pathway. The upregulated genes vary between cohorts and those upregulated in TERT^+^ LMS do not share common Gene Ontology terms or known protein interactions (Fig. S13B). Similarly, pathways upregulated in ALT^−^ pediatric osteosarcomas are not observed in TERT^+^ LMS (17). The genes upregulated in TERT^+^ tumors are more likely to reflect a “private biology” specific to small tumor subsets, varying with tumor histotype. These findings suggest that there is no real biological signature in non-translocation-related sarcomas following telomerase activation. Nonetheless, the number of TERT^+^/C1 tumors in each cohort is limited and further studies with larger sample sizes are needed to confirm these observations. Additionally, no mutations in the *TERT* promoter were detected in LMS or in TERT^+^ hybrids (data not shown), so the mechanism of TERT reactivation in these samples remains unclear. Further investigation into the TERT reactivation processes in sarcomas would be valuable. Several mechanisms have been proposed, including enhancer hijacking due to genomic rearrangements, promoter methylation, or hypermethylation of the repressive locus upstream of the *TERT* promoter (9). Such characterization could help to determine whether distinct subgroups of TERT^+^ tumors exist, which might explain the variability observed between cohorts.

CINSARC is a gene signature associated with chromosomal instability, increased ploidy and high metastatic risk (29,40,41). ALT is also described as related to higher ploidy, promoting chromosomal instability and conferring poor prognosis (2,14,42). Telomere shortening leading to replicative crisis is known to induce cellular stress and subsequent genomic instability through telomere fusions and structural chromosomal anomalies (43,44). Telomere dysfunction can even trigger tetraploidization via defective cell cycle checkpoints (45), which may ultimately generate aneuploidy. These abnormalities can accumulate until cells either die or acquire a telomere maintenance mechanism. WGD and chromosomal instability (CIN) are frequently observed in tumors, particularly in non-translocation-related sarcomas, which have the highest proportion of highly rearranged tumors that activate the ALT mechanism and have undergone a WGD event (7–9,46). The ALT process itself can be a source of genome instability through the insertion of telomeric sequences into multiple internal chromosomal sites (14). The CIN observed in the ALT^+^/CINSARC^high^ tumors may represent the link between ALT/CINSARC and tumor aggressiveness. Bakhoum *et al.* demonstrated that CIN promotes metastasis by generating cytosolic DNA, which activates an immune response mediated by the cGAS-STING pathway (47). This pathway senses the presence of inappropriate DNA in the cytosol, whether from pathogens or from self-DNA arising due to CIN, triggering an immune response. Although initially considered to have anti-tumor effects, increasing evidence suggests that chronic activation of the cGAS-STING pathway in tumors, particularly those with high levels of CIN, can promote tumorigenesis (48). Notably, non-canonical signaling of cGAS-STING triggered by CIN leads to downstream activation of NF-κB pathway (47,49). Tumor cells with elevated CIN may thus exploit chronic innate immune pathway activation to spread to distant organs (47) and might even depend on this non-canonical cGAS-STING-IL6-STAT3 pathway for survival (47,49). The ALT mechanism, which induces CIN and causes accumulation of abundant extrachromosomal telomeric repeats (especially abundant in tumors with high CINSARC enrichment), has been shown to activate immune responses via cGAS-STING activation (50,51), although some prior results suggested that the cGAS-STING pathway may be inactivated in ALT^+^ cells (52). Further studies are needed to precisely evaluate the activation of this pathway in ALT^+^ non-translocation-related sarcomas and to determine whether these tumors depend on IL6 for survival. This could open avenues for testing IL6R inhibitors, potentially enhancing therapeutic efficiency in those chemotherapy-poor-responding tumors. Ultimately, patient management could be improved by identifying tumors likely to benefit from such treatments or even from ALT inhibitors.

Since CINSARC is also a prognostic factor in other cancer types (40), particularly those characterized mainly by telomerase reactivation, it would be insightful to investigate whether CINSARC in these cancers is similarly associated with the ALT phenotype. Such an association would definitely demonstrate the CINSARC/ALT relationship triggering survival and aggressiveness.

## Conclusion

In conclusion, our study shows that in ALT^+^ non-translocation-related sarcomas CINSARC is highly expressed whereas it is expressed at lower level in TERT^+^ tumors. In cellular models, spontaneous activation of ALT is associated with increased CINSARC expression, while inhibition of ALT leads to decreased CINSARC expression. Altogether, our data support the concept that CINSARC is intrinsically linked to ALT in non-translocation-related sarcomas. Nevertheless, additional studies are required to precisely delineate the roles of individual CINSARC genes in ALT and to explore this link in other tumor types.

## Abbreviations

ALT: Alternative Lengthening of Telomeres
APB: ALT-associated PML bodies
CIN: Chromosomal Instability
CINSARC: Complexity Index in Sarcomas
DEG: Differentially Expressed Genes
GO: Gene Ontology
GSEA: Gene Set Enrichment Analysis
ICGC: International Cancer Genome Consortium
LMS: Leiomyosarcoma
NES: Normalized Enrichment Score
RNAseq: RNA sequencing
SD: Standard Deviation
TERT: Telomerase
TMM: Telomere Maintenance Mechanism
WGD: Whole Genome Doubling
WGseq: Whole Genome Sequencing.

## DECLARATIONS

### Ethics Statement

The ICGC study was approved by the Ethical Committee of the Sud-Ouest et Outre Mer III Committee for the Protection of Individuals (DC 2014/47 bis), and written informed consent was obtained from all patients.

For cohort 2, tumor tissue samples were obtained from patients and stored at the Centre de Ressources Biologiques de l’Institut Bergonié (CRB-IB). At the time of collection, patients provided written informed consent or expressed non-opposition for the use of their biological samples for research purposes. The RNA sequencing study for cohort 2 was reviewed and approved by the Institut Bergonié Ethical Committee.

All samples were part of the CRB-IB, which is authorized by the French authorities to collect and use human biological samples for research, in accordance with the French Public Health Code (articles L.1243-4 and R.1243-61).

Myoblast cells from a healthy donor were collected after written informed consent, with protocol approval from the Montpellier University Hospitals ethics committee (2009-04-BPCO-V2).

All methods were conducted in accordance with approved protocols, the French Public Health Code, applicable national regulations, and the principles of the Declaration of Helsinki.

### Consent for publication

Not applicable

### Data Availability Statement

**Cohort 1**: ICGC RNAseq fastq files for the 98 LMS will be available at the time of publication at https://ega-archive.org/datasets/ under dataset ID: EGAD00001003192: https://ega-archive.org/datasets/EGAD00001003192.

**Cohort 2**: RNAseq fastq files for the 99 of the 106 RNAseq are available on the NCBI Sequence Read Archive (SRA) (RRID:SCR_004891) under accession number: SRP057793. The 7 remaining will be available at the time of publication on SRA under accession number: PRJNA1331653.

**Cohort 3**: TCGA RNAseq data were downloaded from the National Cancer Institute GDC Data Portal: https://portal.gdc.cancer.gov/.

**Hybrid cell lines and *BLM* shRNA models:** RNAseq data will be available on SRA under accession number at the time of publication: PRJNA1314156.

### Competing interests

The authors declare that they have no competing interests.

### Funding

Not applicable

### Author Contributions

**GP:** Conceptualization, data curation, formal analysis, investigation, methodology, validation visualization, writing-original draft, writing-review and editing. **CG**: Formal analysis, investigation. **NR**: Data curation, software, formal analysis, investigation. **ES**: Formal analysis, investigation. **CV**: Formal analysis, investigation. **PP**: Formal analysis, investigation, validation. **FT**: Formal analysis, software, investigation. **DP**: Data curation, formal analysis, investigation, methodology, validation. **FC**: Conceptualization, formal analysis, software, funding acquisition, methodology, project administration, supervision, visualization, writing–original draft, writing–review and editing. All authors read and approved the final manuscript.

## Acknowledgments

Part of Bioinformatics analyses were performed thanks to the Core Cluster of the Genotoul (Genopole Toulouse). The results shown here are partly based upon data generated by the TCGA Research Network: https://www.cancer.gov/ccg/access-data. We are thankful to the French Sarcoma Group for tumor banks. We are grateful to the INCa (Institute National du Cancer), the *Phil’anthrope* and the *Pour Corentin* associations for their financial support.

## Notes

### Competing Interest Statement

The authors have declared no competing interest.

## References

1. Hoang SM, O’Sullivan RJ. Alternative Lengthening of Telomeres: Building Bridges To Connect Chromosome Ends. Trends Cancer. 2020;6:247–60.

2. Dilley RL, Greenberg RA. ALTernative Telomere Maintenance and Cancer. Trends Cancer. 2015;1:145–56.

3. Zhang JM, Zou L. Alternative lengthening of telomeres: from molecular mechanisms to therapeutic outlooks. Cell Biosci. 2020;10:30.

4. Kockler ZW, Comeron JM, Malkova A. A unified alternative telomere-lengthening pathway in yeast survivor cells. Mol Cell. 2021;81:1816–1829.e5.

5. Chang S, Tan J, Bao R, Zhang Y, Tong J, Jia T, et al. Multiple functions of the ALT favorite helicase, BLM. Cell Biosci. 2025;15:31.

6. WHO Classification of Tumours Editorial Board. Soft Tissue and Bone Tumours. 5th ed. Vol. 3. Lyon (France): International Agency for Research on Cancer: IARC press; 2020.

7. Cancer Genome Atlas Research Network. Comprehensive and Integrated Genomic Characterization of Adult Soft Tissue Sarcomas. Cell. 2017;171:950–965.e28.

8. Chudasama P, Mughal SS, Sanders MA, Hübschmann D, Chung I, Deeg KI, et al. Integrative genomic and transcriptomic analysis of leiomyosarcoma. Nat Commun. 2018;9:144.

9. Steele CD, Tarabichi M, Oukrif D, Webster AP, Ye H, Fittall M, et al. Undifferentiated Sarcomas Develop through Distinct Evolutionary Pathways. Cancer Cell. 2019;35:441–456.e8.

10. Costa A, Daidone MG, Daprai L, Villa R, Cantù S, Pilotti S, et al. Telomere Maintenance Mechanisms in Liposarcomas: Association with Histologic Subtypes and Disease Progression. Cancer Res. 2006;66:8918–24.

11. Henson JD, Hannay JA, McCarthy SW, Royds JA, Yeager TR, Robinson RA, et al. A Robust Assay for Alternative Lengthening of Telomeres in Tumors Shows the Significance of Alternative Lengthening of Telomeres in Sarcomas and Astrocytomas. Clin Cancer Res. 2005;11:217–25.

12. Liau JY, Lee JC, Tsai JH, Yang CY, Liu TL, Ke ZL, et al. Comprehensive screening of alternative lengthening of telomeres phenotype and loss of ATRX expression in sarcomas. Mod Pathol. 2015;28:1545–54.

13. Ballinger ML, Pattnaik S, Mundra PA, Zaheed M, Rath E, Priestley P, et al. Heritable defects in telomere and mitotic function selectively predispose to sarcomas. Science. 2023;379:253–60.

14. Marzec P, Armenise C, Pérot G, Roumelioti FM, Basyuk E, Gagos S, et al. Nuclear-Receptor-Mediated Telomere Insertion Leads to Genome Instability in ALT Cancers. Cell. 2015;160:913–27.

15. Lawlor RT, Veronese N, Pea A, Nottegar A, Smith L, Pilati C, et al. Alternative lengthening of telomeres (ALT) influences survival in soft tissue sarcomas: a systematic review with meta-analysis. BMC Cancer. 2019;19:232.

16. Mori JO, Keegan J, Flynn RL, Heaphy CM. Alternative lengthening of telomeres: mechanism and the pathogenesis of cancer. J Clin Pathol. 2024;77:82–6.

17. De Nonneville A, Salas S, Bertucci F, Sobinoff AP, Adélaïde J, Guille A, et al. TOP3A amplification and ATRX inactivation are mutually exclusive events in pediatric osteosarcomas using ALT. EMBO Mol Med. 2022;14:e15859.

18. Darmusey L, Pérot G, Thébault N, Le Guellec S, Desplat N, Gaston L, et al. ATRX Alteration Contributes to Tumor Growth and Immune Escape in Pleomorphic Sarcomas. Cancers. 2021;13:2151.

19. Lesluyes T, Pérot G, Largeau MR, Brulard C, Lagarde P, Dapremont V, et al. RNA sequencing validation of the Complexity INdex in SARComas prognostic signature. Eur J Cancer. 2016;57:104–11.

20. Pomiès P, Rodriguez J, Blaquière M, Sedraoui S, Gouzi F, Carnac G, et al. Reduced myotube diameter, atrophic signalling and elevated oxidative stress in cultured satellite cells from COPD patients. J Cell Mol Med. 2015;19:175–86.

21. Dobin A, Davis CA, Schlesinger F, Drenkow J, Zaleski C, Jha S, et al. STAR: ultrafast universal RNA-seq aligner. Bioinformatics. 2013;29:15–21.

22. Danecek P, Bonfield JK, Liddle J, Marshall J, Ohan V, Pollard MO, et al. Twelve years of SAMtools and BCFtools. GigaScience. 2021;10:giab008.

23. Putri GH, Anders S, Pyl PT, Pimanda JE, Zanini F. Analysing high-throughput sequencing data in Python with HTSeq 2.0. Boeva V, editor. Bioinformatics. 2022;38:2943–5.

24. Love MI, Huber W, Anders S. Moderated estimation of fold change and dispersion for RNA-seq data with DESeq2. Genome Biol. 2014;15:550.

25. Subramanian A, Tamayo P, Mootha VK, Mukherjee S, Ebert BL, Gillette MA, et al. Gene set enrichment analysis: A knowledge-based approach for interpreting genome-wide expression profiles. Proc Natl Acad Sci U A. 2005;102:15545–50.

26. Déjardin J, Kingston RE. Purification of Proteins Associated with Specific Genomic Loci. Cell. 2009;136:175–86.

27. Dal Col P, Poncet D, Rivoirard R, Vassal F, Bernichon E, Boutet C, et al. Acquired ATRX Loss and ALT Phenotype Through Tumor Recurrences in a Case of Pleomorphic Xanthoastrocytoma Suggest Their Possible Roles in Tumor Progression. J Neuropathol Exp Neurol. 2020;79:1011–4.

28. Pérot G, Derré J, Coindre JM, Tirode F, Lucchesi C, Mariani O, et al. Strong Smooth Muscle Differentiation Is Dependent on Myocardin Gene Amplification in Most Human Retroperitoneal Leiomyosarcomas. Cancer Res. 2009;69:2269–78.

29. Chibon F, Lagarde P, Salas S, Pérot G, Brouste V, Tirode F, et al. Validated prediction of clinical outcome in sarcomas and multiple types of cancer on the basis of a gene expression signature related to genome complexity. Nat Med. 2010;16:781–7.

30. Szklarczyk D, Gable AL, Lyon D, Junge A, Wyder S, Huerta-Cepas J, et al. STRING v11: protein–protein association networks with increased coverage, supporting functional discovery in genome-wide experimental datasets. Nucleic Acids Res. 2019;47:D607–13.

31. Lartigue L. Genome remodeling upon mesenchymal tumor cell fusion contributes to tumor progression and metastatic spread. :14.

32. Delespaul L, Merle C, Lesluyes T, Lagarde P, Guellec SL, Pérot G, et al. Fusion-mediated chromosomal instability promotes aneuploidy patterns that resemble human tumors. Oncogene. 2019 July 3;1–12.

33. Delespaul L, Gélabert C, Lesluyes T, Le Guellec S, Pérot G, Leroy L, et al. Cell–cell fusion of mesenchymal cells with distinct differentiations triggers genomic and transcriptomic remodelling toward tumour aggressiveness. Sci Rep. 2020 Dec;10(1):21634.

34. Kaur E, Agrawal R, Sengupta S. Functions of BLM Helicase in Cells: Is It Acting Like a Double-Edged Sword? Front Genet. 2021;12:634789.

35. Pan X, Drosopoulos WC, Sethi L, Madireddy A, Schildkraut CL, Zhang D. FANCM, BRCA1, and BLM cooperatively resolve the replication stress at the ALT telomeres. Proc Natl Acad Sci. 2017;114:E5940–9.

36. Shi G, Hu Y, Zhu X, Jiang Y, Pang J, Wang C, et al. A critical role of telomere chromatin compaction in ALT tumor cell growth. Nucleic Acids Res. 2020;48:6019–31.

37. Kaminski N, Wondisford AR, Kwon Y, Lynskey ML, Bhargava R, Barroso-González J, et al. RAD51AP1 regulates ALT-HDR through chromatin-directed homeostasis of TERRA. Mol Cell. 2022;82:4001–4017.e7.

38. Bhattacharyya S, Keirsey J, Russell B, Kavecansky J, Lillard-Wetherell K, Tahmaseb K, et al. Telomerase-associated Protein 1, HSP90, and Topoisomerase IIα Associate Directly with the BLM Helicase in Immortalized Cells Using ALT and Modulate Its Helicase Activity Using Telomeric DNA Substrates. J Biol Chem. 2009;284:14966–77.

39. Jemaà M, Abdallah S, Lledo G, Perrot G, Lesluyes T, Teyssier C, et al. Heterogeneity in sarcoma cell lines reveals enhanced motility of tetraploid versus diploid cells. Oncotarget. 2017;8:16669–89.

40. Chibon F, Lesluyes T, Valentin T, Le Guellec S. CINSARC signature as a prognostic marker for clinical outcome in sarcomas and beyond. Genes Chromosomes Cancer. 2019;58:124–9.

41. Lesluyes T, Chibon F. A Global and Integrated Analysis of CINSARC-Associated Genetic Defects. Cancer Res. 2020;80:5282–90.

42. Christodoulidou A, Raftopoulou C, Chiourea M, Papaioannou GK, Hoshiyama H, Wright WE, et al. The Roles of Telomerase in the Generation of Polyploidy during Neoplastic Cell Growth. Neoplasia. 2013;15:156–IN17.

43. Zou Y, Sfeir A, Gryaznov SM, Shay JW, Wright WE. Does a Sentinel or a Subset of Short Telomeres Determine Replicative Senescence? Mol Biol Cell. 2004;15:3709–18.

44. Cleal K, Baird DM. Catastrophic Endgames: Emerging Mechanisms of Telomere-Driven Genomic Instability. Trends Genet. 2020;36:347–59.

45. Davoli T, Denchi EL, De Lange T. Persistent Telomere Damage Induces Bypass of Mitosis and Tetraploidy. Cell. 2010;141:81–93.

46. Beird HC, Wu CC, Nakazawa M, Ingram D, Daniele JR, Lazcano R, et al. Complete loss of TP53 and RB1 is associated with complex genome and low immune infiltrate in pleomorphic rhabdomyosarcoma. Hum Genet Genomics Adv. 2023;4:100224.

47. Bakhoum SF, Ngo B, Laughney AM, Cavallo JA, Murphy CJ, Ly P, et al. Chromosomal instability drives metastasis through a cytosolic DNA response. Nature. 2018;553:467–72.

48. Lanng KRB, Lauridsen EL, Jakobsen MR. The balance of STING signaling orchestrates immunity in cancer. Nat Immunol. 2024;25:1144–57.

49. Hong C, Schubert M, Tijhuis AE, Requesens M, Roorda M, Van Den Brink A, et al. cGAS–STING drives the IL-6-dependent survival of chromosomally instable cancers. Nature. 2022;607:366–73.

50. Segura-Bayona S, Villamor-Payà M, Attolini CSO, Koenig LM, Sanchiz-Calvo M, Boulton SJ, et al. Tousled-Like Kinases Suppress Innate Immune Signaling Triggered by Alternative Lengthening of Telomeres. Cell Rep. 2020;32:107983.

51. Emam A, Wu X, Xu S, Wang L, Liu S, Wang B. Stalled replication fork protection limits cGAS–STING and P-body-dependent innate immune signalling. Nat Cell Biol. 2022;24:1154–64.

52. Chen YA, Shen YL, Hsia HY, Tiang YP, Sung TL, Chen LY. Extrachromosomal telomere repeat DNA is linked to ALT development via cGAS-STING DNA sensing pathway. Nat Struct Mol Biol. 2017;24:1124–31.

