## Supplemental Figures for "Alternative Lengthening of Telomeres and CINSARC are interconnected toward non-translocation-related sarcomas progression"

### Supplementary Figures and legends

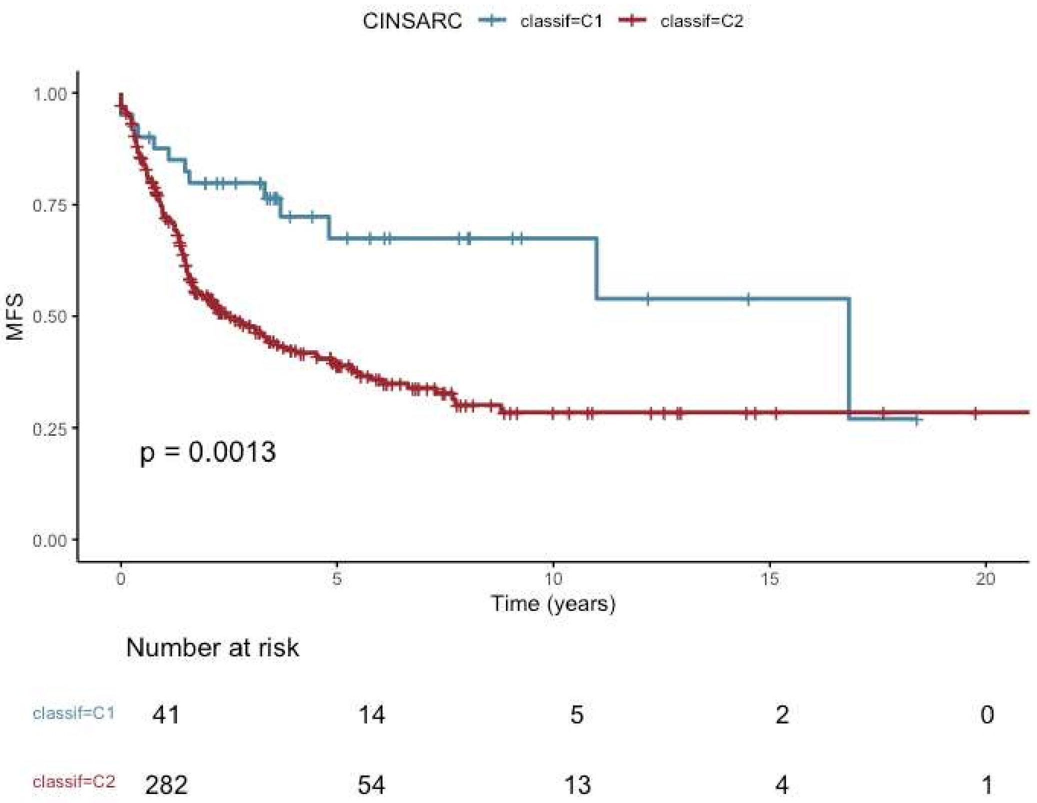


**Figure S1: CINSARC prognostic value in the three grouped cohorts.**

Kaplan-Meier mestastasis-free survival (MFS) analysis in non-translocation-related sarcomas of the three cohorts classified according to CINSARC groups. The C1 classification corresponds to tumors with no CINSARC enrichment, while the C2 classification corresponds to tumors with a significant positive CINSARC enrichment score. Ninety-seven patients of cohort 1 (1 not significant case, Table S1) were grouped with the 106 patients of cohort 2 and the 120 patients of cohort 3 (n=323). Y-axis indicates the Log-Rank test p-value. The number at risk indicates, for each CINSARC group, the number of subjects who have not experienced the event of interest (MFS) or been censored at every time point (every 5 years).

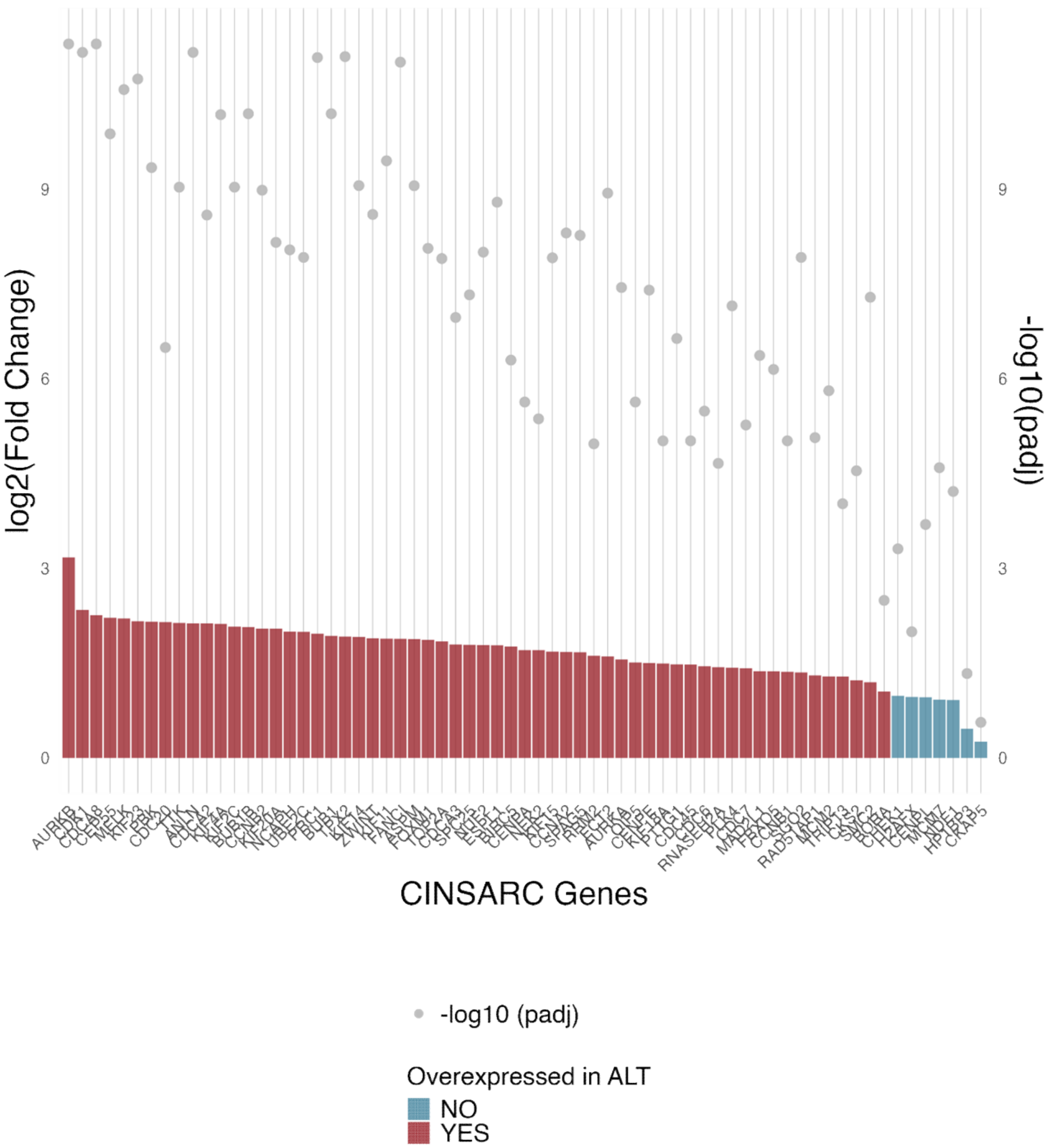


**Figure S2: CINSARC genes upregulation in ALT⁺ LMS.**

Graphical representation of log2FC and adj. p-value (Benjamini-Hochberg correction) obtained for the 67 genes of the CINSARC signature from the DEG analysis in LMS. Comparison between ALT$^+$ LMS versus TERT$^+$ LMS. Significantly upregulated CINSARC genes in ALT$^+$ LMS are indicated in red on the bar chart (adj. p-value<0.05 and log2FC>1).

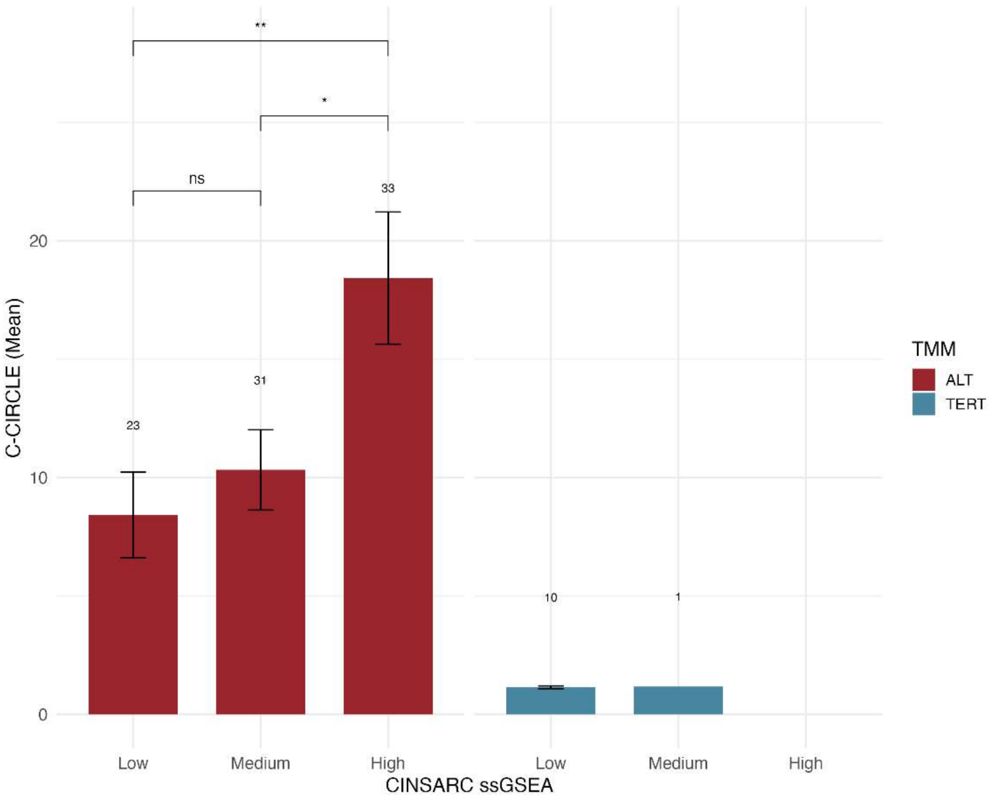


**Figure S3: The more CINSARC is enriched in tumors, the more C-circles they have.**

The 98 LMS from the cohort 1 were separated in three equal groups according to their CINSARC ssGSEA NES (low, medium and high NES). The bar plot presents the mean of C-circles in those three groups with a distinction between $ALT^+$ and $TERT^+$ tumors. Number of cases in each group is indicated above each bar. Mann-Whitney test. ns: not significant, *: $p < 0.05$, **: $p < 0.01$.

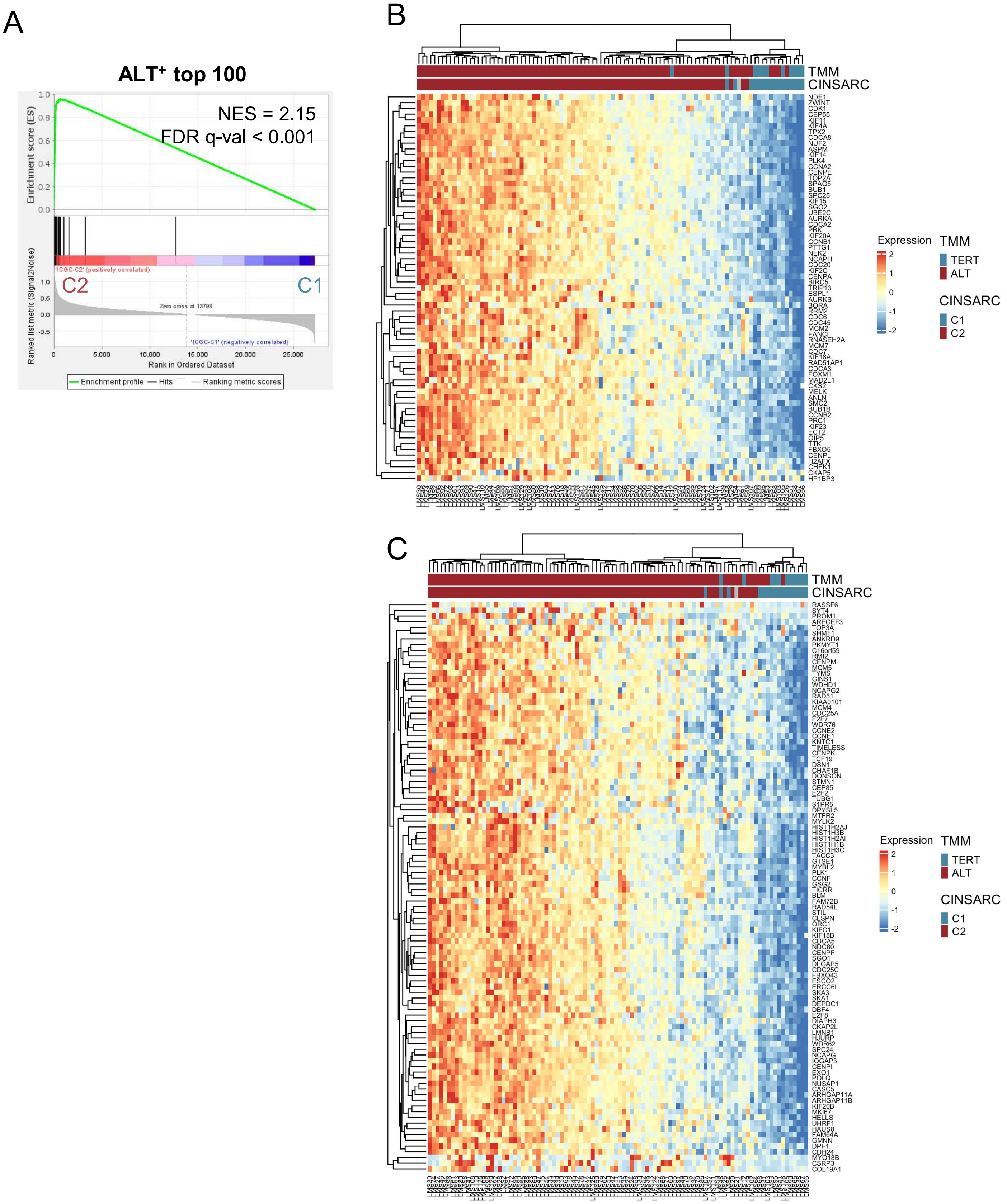


**Figure S4: CINSARC and ALT⁺-related signatures classify LMS similarly (ICGC cohort).**

**A.** Enrichment plot of the ALT⁺-related gene signature (100 most significantly upregulated coding genes in ALT⁺ LMS) in the two CINSARC tumor groups (C1: tumors with a low CINSARC enrichment score and C2: tumors with a high CINSARC enrichment score). **B.** Unsupervised hierarchical clustering heatmap of the 98 LMS from ICGC cohort using the 67 genes of the CINSARC signature. CINSARC groups and telomere maintenance mechanism (TMM) are indicated. **C.** Unsupervised hierarchical clustering heatmap of the 98 LMS from ICGC cohort using the ALT⁺-related signature. CINSARC groups and TMM are indicated.

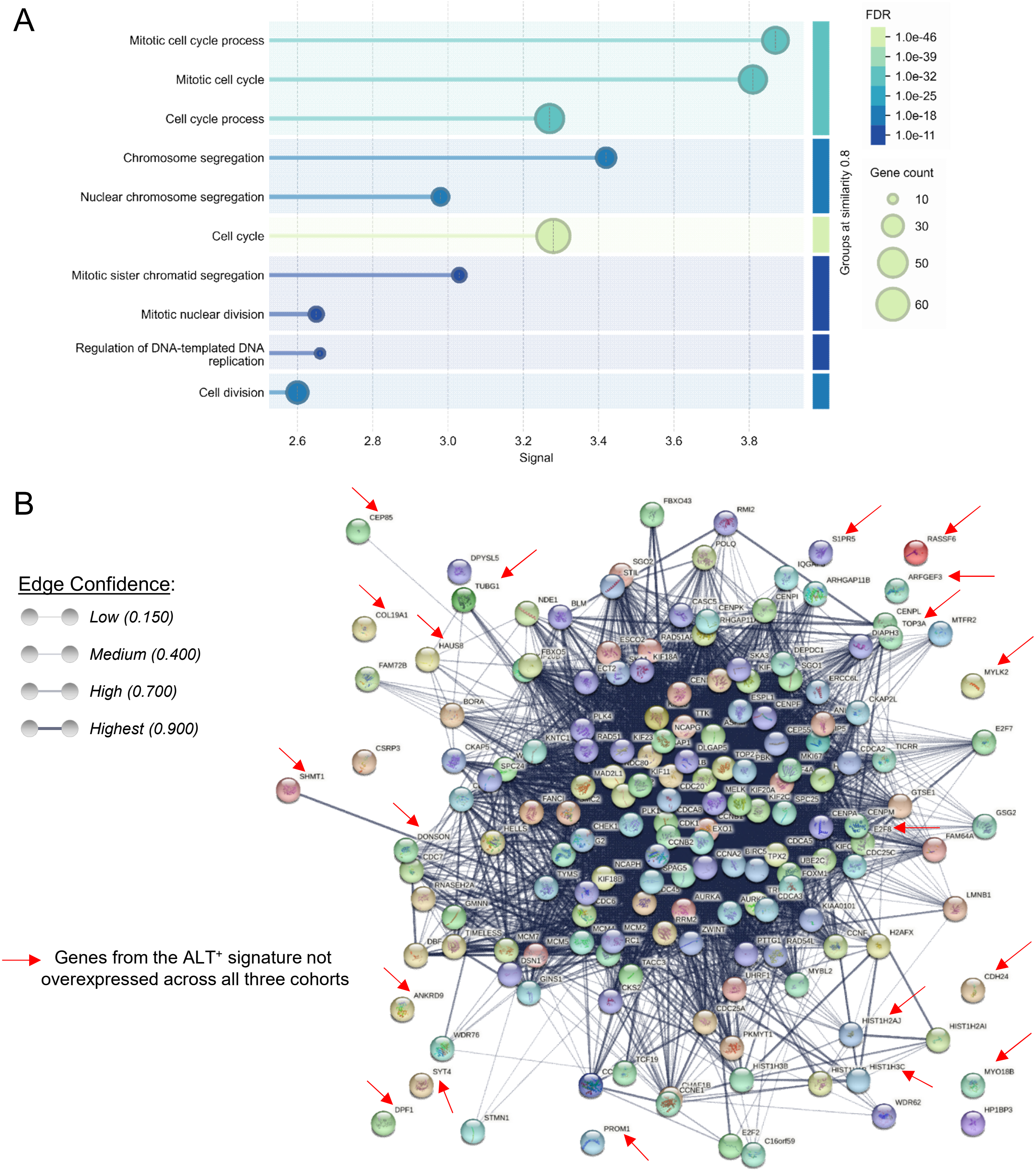


**Figure S5: Pathways involving the 100 genes of the ALT⁺-related signature are the same as those for the CINSARC genes.**

**A.** Lollipop plot representation of the Gene Ontology analysis of the network of the 100 most upregulated coding genes in ALT⁺ LMS performed on STRING database website (v12.0). Only the 10 most significant biological processes are represented. **B.** STRING network of proteins encoded by the 100 genes of ALT⁺-related and the 67 genes of CINSARC signatures obtained from STRING database (v12.0). Full STRING network with confidence edges. Active interaction sources: experiments, databases and co-expression. Number of nodes: 167 (nodes represent proteins), number of edges: 4924 (edges represent protein-protein functional and physical associations), average node degree: 59, avg. local clustering coefficient: 0.704, expected number of edges: 336, PPI enrichment p-value: $< 10^{-16}$. Line thickness indicates the strength of data support (edge confidence).

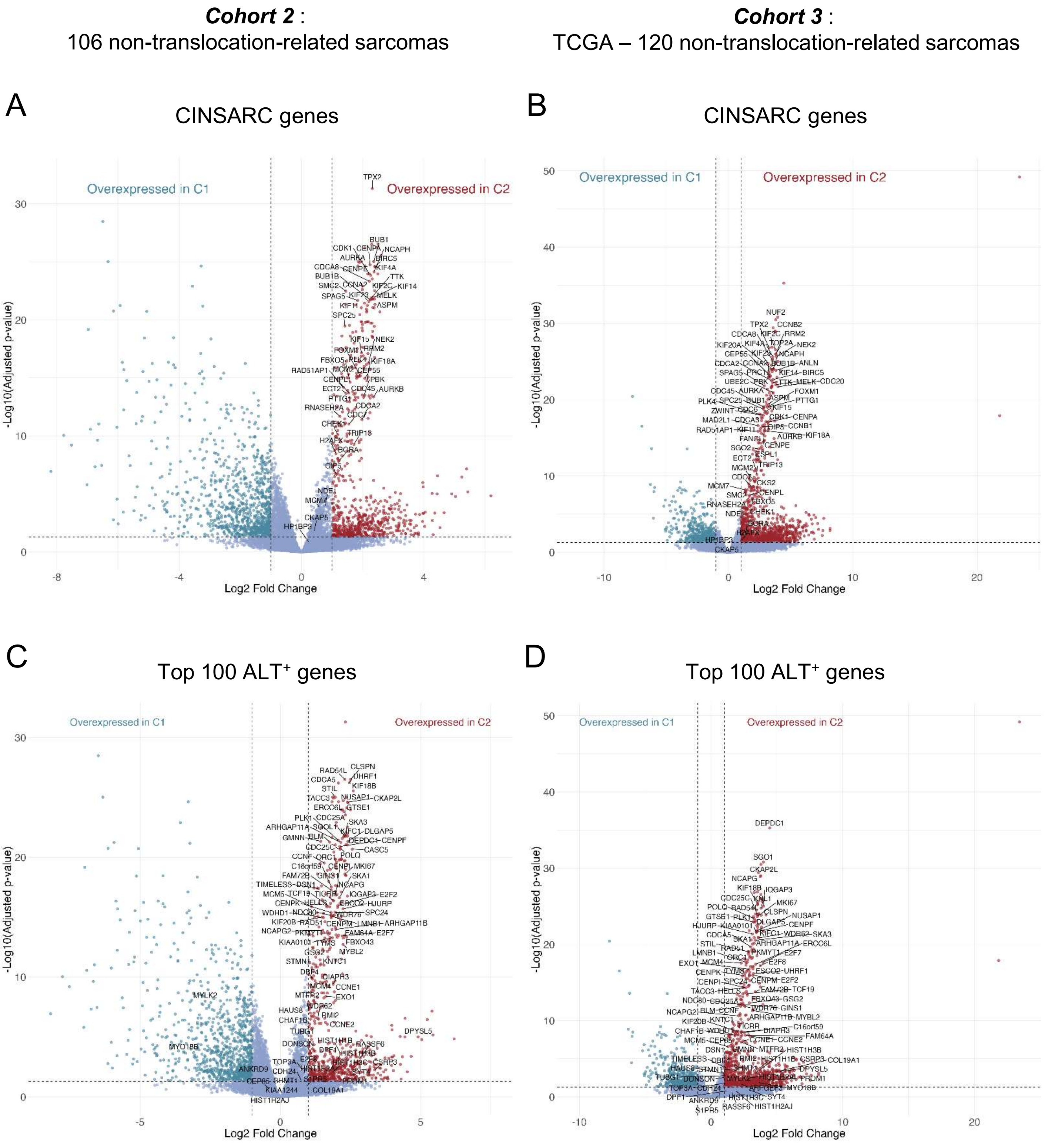


**Figure S6: CINSARC and ALT⁺-related genes behave similarly in all cohorts.**

Volcano plots showing the differentially expressed genes (DEG) between C2 and C1 tumors from cohort 2 (**A and C**) or from TCGA cohort (**B and D**). C2 tumors are those enriched for CINSARC gene signature and C1 tumors those not enriched. Dotted lines highlight thresholds used to define DEG: adj. p-value < 0.05 (Benjamini-Hochberg correction) and absolute log2Fold change > 1. **A and B.** The 67 CINSARC genes are annotated on the 2 volcano plots. **C and D**. The 100 genes from the ALT⁺-related signature are annotated on the 2 volcano plots.

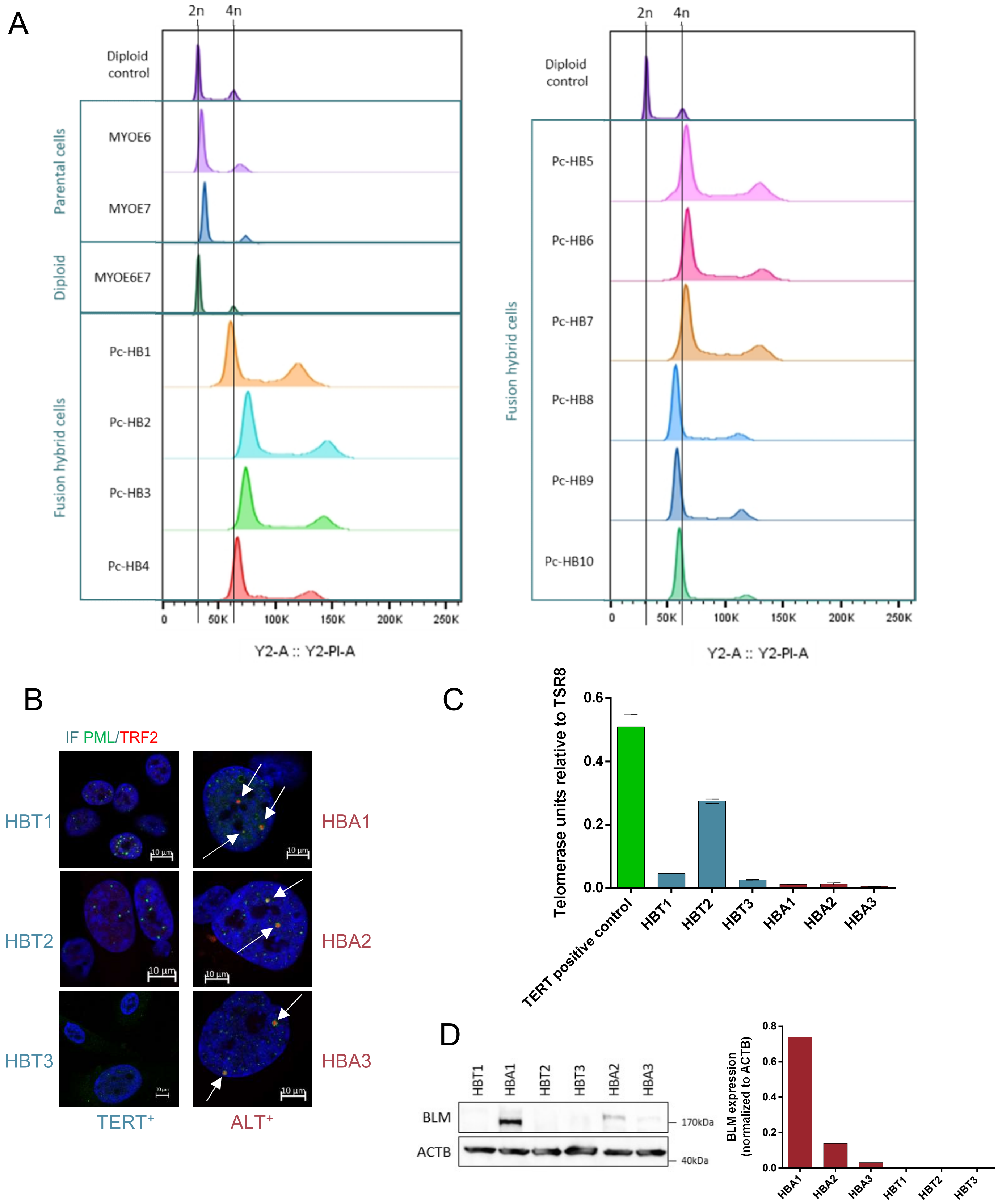


**Figure S7: Validation of the cellular models.**

**A.** Ploidy assessment of the cell lines using Propidium iodide labelling. **B.** PML/TERF2 co-immunofluorescence. Pictures obtained from confocal analysis show a representative PML (green) and TERF2 (red) labeling in nuclei (labeled with Dapi) for each cell line. Arrows highlight the TERF2/PML co-localized signals corresponding to ALT-associated PML bodies (APB). Scale bar: 10 µm. **C.** TRAPeze® RT Telomerase assay: TERT units, relative to TSR8, in each cell line. **D.** BLM expression in hybrid cell lines by western blot. The bar plot presents the quantification of BLM expression, after normalization to the b-actin expression (ACTB), for each sample. Pc-HB: pre-crisis hybrid. HBT: hybrids TERT$^{+}$ , HBA: hybrids ALT$^{+}$. Uncropped blot for BLM is presented in Supplementary Figure S14.

A

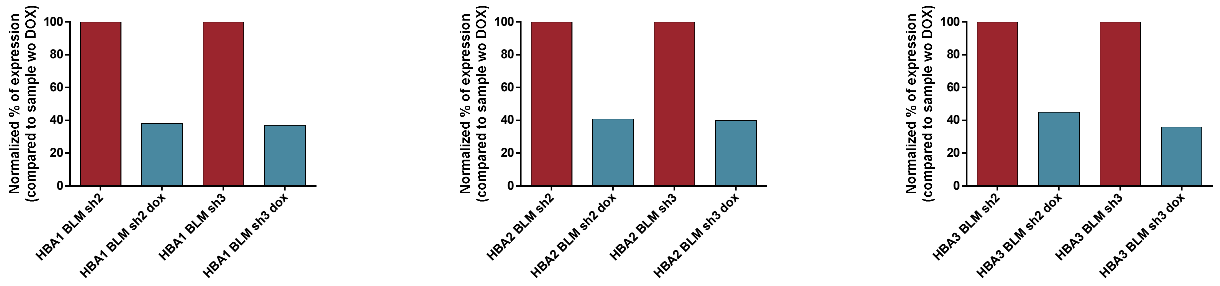


B

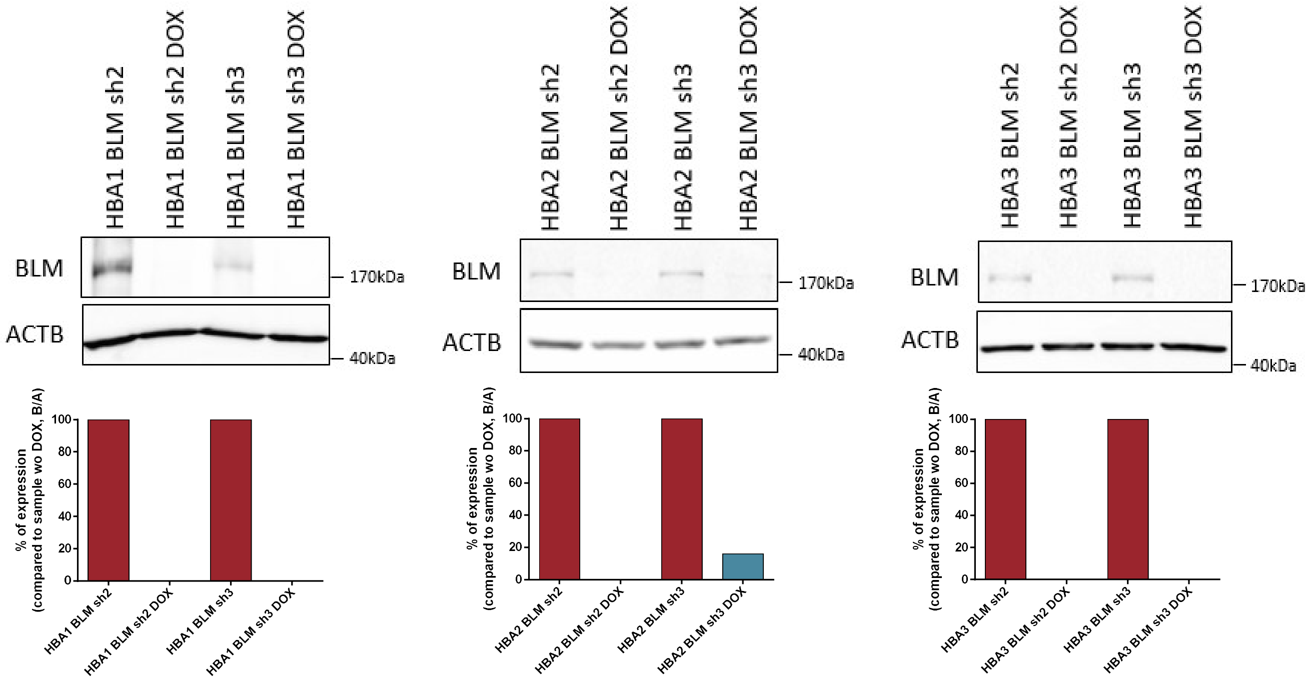


**Figure S8: Validation of the cellular models.**

**A.** Real-time quantitative RT-PCR after 4 weeks of DOX treatment of three ALT$^{+}$ cell lines. For each sample, *BLM* expression was first normalized to the mean of *ACTB* and *RPLP0* expressions. Bar plots presented the normalized quantification of *BLM* expression for each sample under DOX treatment (*BLM* sh DOX, in blue) compared to the expression of the same cell line without DOX treatment (*BLM* sh, in red). Two different shRNAs were used in each cell line (sh2 & sh3). **B.** Western blot after 4 weeks of DOX treatment. BLM expression was first normalized to the b-actin (ACTB) for each sample. Bar plots presented the normalized quantification of BLM expression for each sample under DOX treatment compared to the expression of the same cell line without DOX treatment. Uncropped blots for BLM are presented in Supplementary Figure S15.

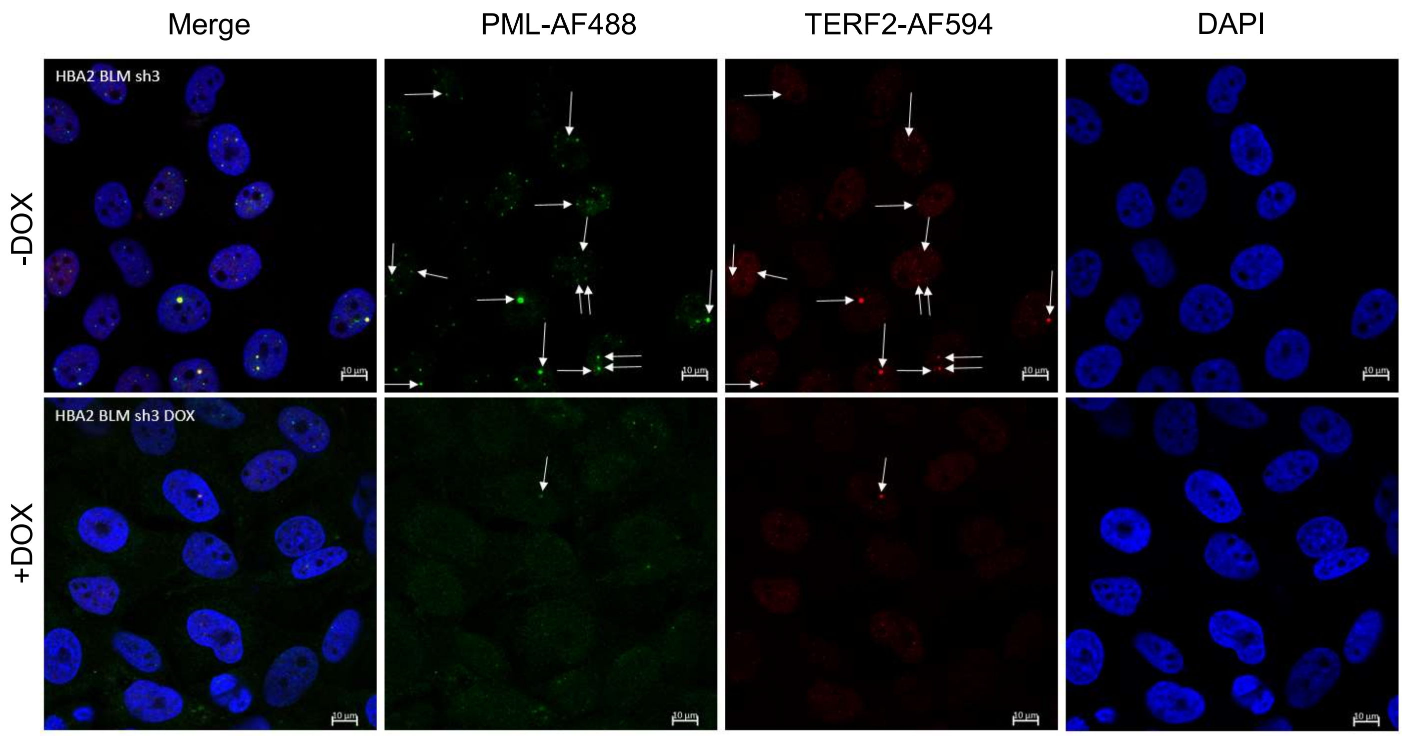


**Figure S9:** ***BLM*** **inhibition reduces the number of APB⁺ cells in the three ALT⁺ cell lines.**

PML/TERF2 co-immunofluorescence results on ALT⁺ cell lines with or without *BLM* inhibition induction. Representative pictures obtained by confocal microscopy for one cell line (HBA2 BLM sh3 ± DOX, treatment of 72h) during one experiment are presented. Arrows highlight the TERF2/PML co-localized signals corresponding to APB. Scale bar: 10 μm.

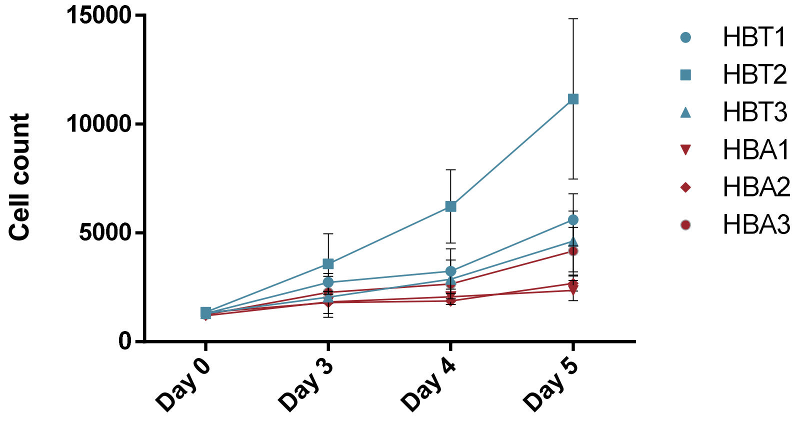


**Figure S10:** ***Proliferation assay of hybrid cell lines*.**

The graph shows the number of cells in the three TERT⁺ cell lines (in blue) and the three ALT⁺ cell lines (in red) over five days. Results are mean of three independent experiments ± SD.

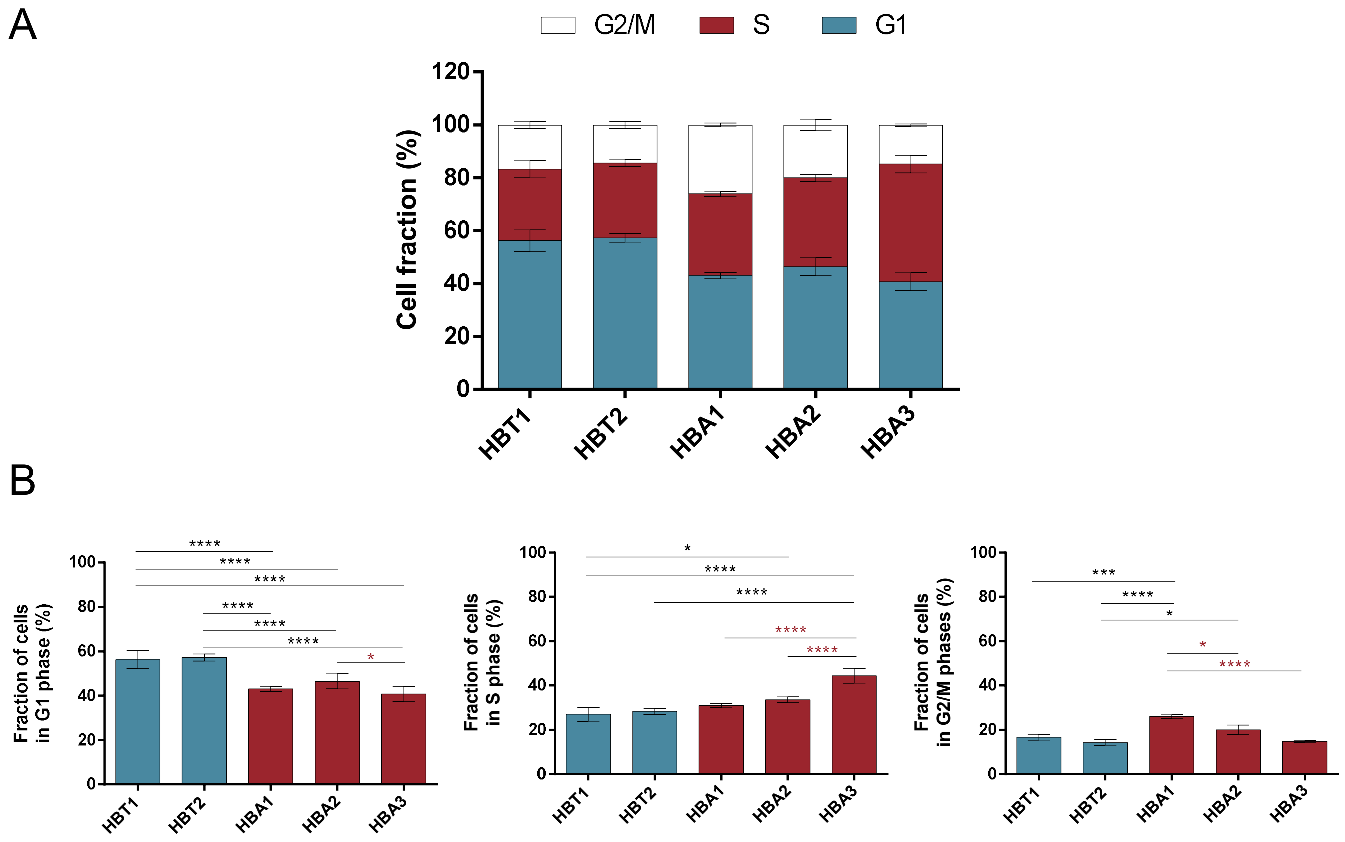


**Figure S11: Hybrid cells repartition during the cell cycle.**

**A.** Cell fraction in G1, S and G2/M phases of the cell cycle assessed by propidium iodide labelling for five hybrid cell lines. HBT3 TERT⁺ cell line could not be analyzed because of the presence of two populations with different ploidies. Results correspond to the mean of three independent experiments ± SD. **B.** Statistical test results: one for each phase of the cell cycle. Two-way ANOVA with Sidak's correction for multiple comparisons test. $TERT^+$ cell lines are indicated in blue and $ALT^+$ ones in red. *: $p < 0.05$, ***: $p < 0.001$ and ****: $p < 0.0001$. * in red show the significant result between 2 $ALT^+$ cell lines.

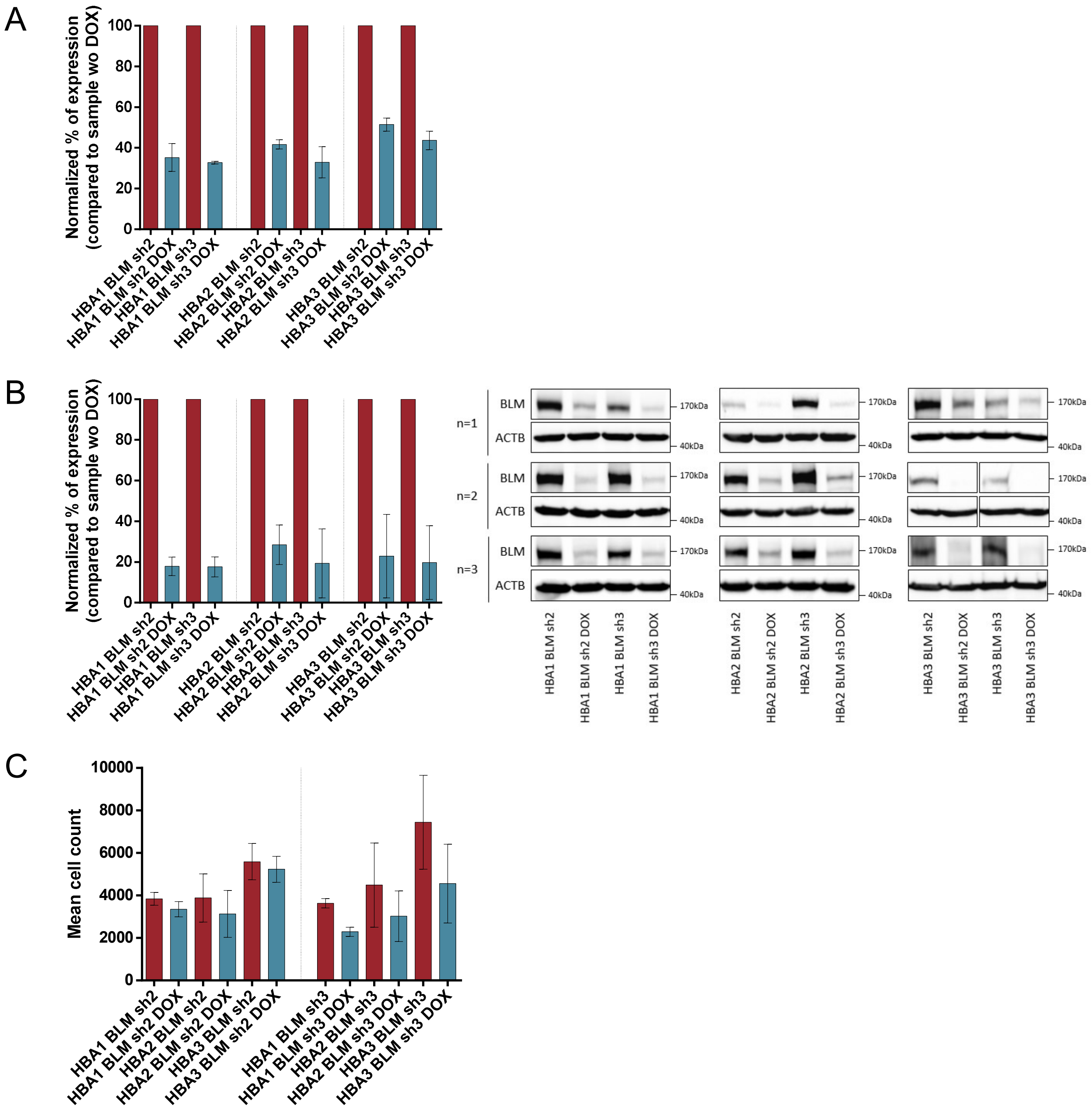


**Figure S12: BLM inhibition validation and cell proliferation assay.**

BLM inhibition validation and proliferation assay for the three ALT$^+$ cell lines with inducible BLM shRNAs after 5 days of DOX treatment. Two different shRNAs were used in each cell line (sh2 & sh3).

**A.** Real-time quantitative RT-PCR validation of *BLM* shRNAS efficiency. For each sample, *BLM* expression was first normalized to the mean of *ACTB* and *RPLP0* expressions. Bar plot presents the normalized quantification of *BLM* expression for each sample under *BLM* inhibition (*BLM* sh DOX, in blue) compared to the expression of the same cell line without DOX treatment (Ctrl, *BLM* sh, in red). Results are presented as the mean of three independent experiments ± SD. **B.** Western blot validation of *BLM* shRNAs efficiency. BLM expression was first normalized to the b-actin (ACTB) for each sample. Bar plot presents the normalized quantification of BLM expression for each sample under BLM inhibition (in blue) compared to the expression of the corresponding control cell line (in red). Results are presented as the mean of three independent experiments ± SD. Uncropped BLM WB are presented in Supplementary Figures S16 and S17. **C.** The bar chart presents the result of the proliferation assay. Results are presented as the mean of three independent experiments ± SD. Mann-Whitney test. No significant result.

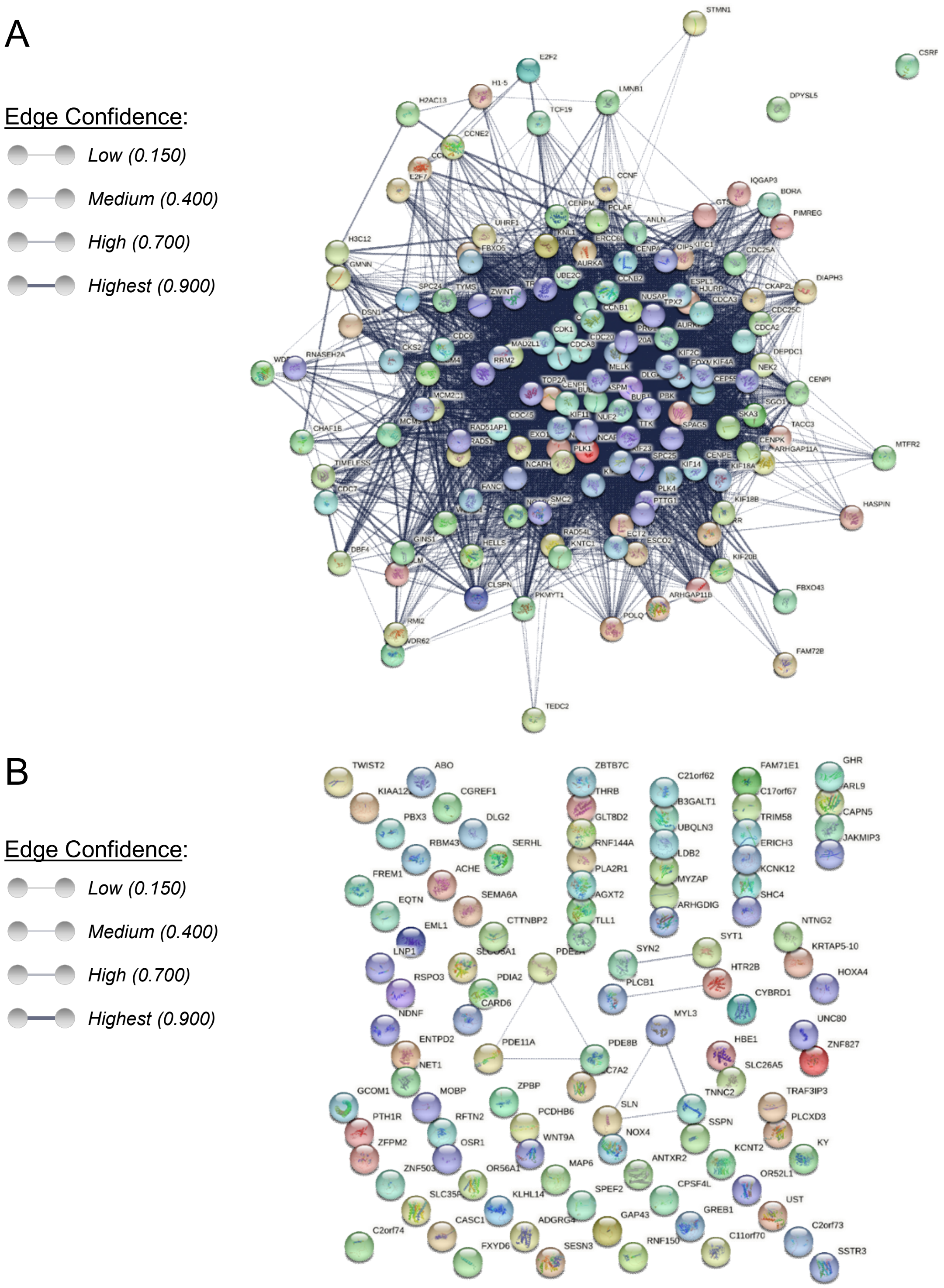


**Figure S13: Networks of proteins encoded by a selection of genes upregulated in ALT$^+$/C2 or TERT$^+$ tumors.**

**A.** STRING network of proteins encoded by the 80 genes of the ALT$^+$-related signature and the 59 CINSARC genes upregulated in ALT$^+$ and C2 tumors across the three cohorts: representation obtained from STRING database (v12.0). Full STRING network with confidence edges. Active interaction sources: experiments, databases and co-expression. Number of nodes: 139 (nodes represent proteins), number of edges: 4542 (edges represent protein-protein functional and physical associations), average node degree: 65.4, avg. local clustering coefficient: 0.771, expected number of edges: 270, PPI enrichment p-value: $< 10^{-16}$. Line thickness indicates the strength of data support (edge confidence). **B.** STRING network of proteins encoded by the 100 coding genes the most significantly upregulated in TERT$^+$ LMS. Representation obtained from STRING database (v12.0). Same parameters as previously were used. Number of nodes: 100, number of edges: 8, average node degree: 17.2, avg. local clustering coefficient: 0.1, expected number of edges: 2, PPI enrichment p-value = 0.00246. Line thickness indicates the strength of data support (edge confidence).

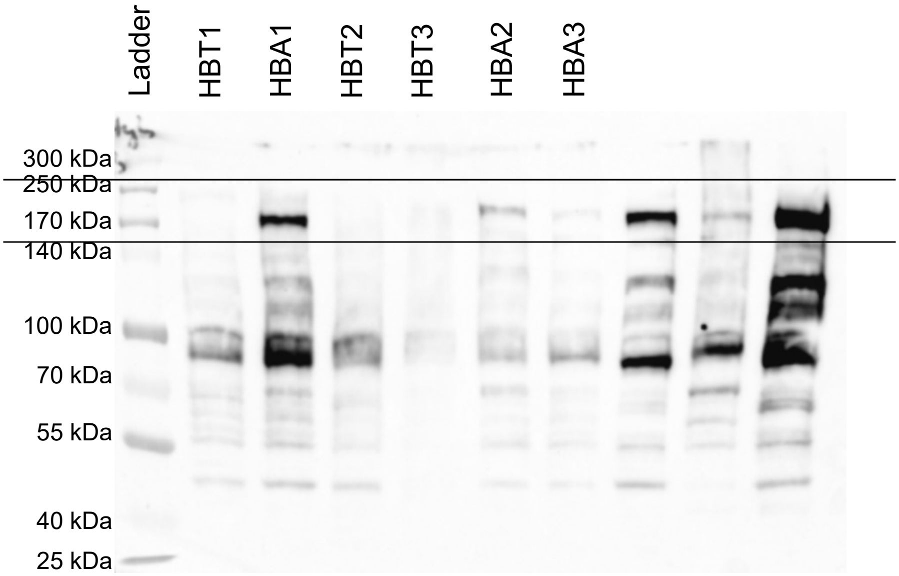


**Figure S14: Uncropped BLM western blot corresponding to Figure S7 D.**

BLM expression in hybrid cell lines by western blot. **The** Ladder used is the ProSieve® QuadColor® marker, Lonza (#00193837).

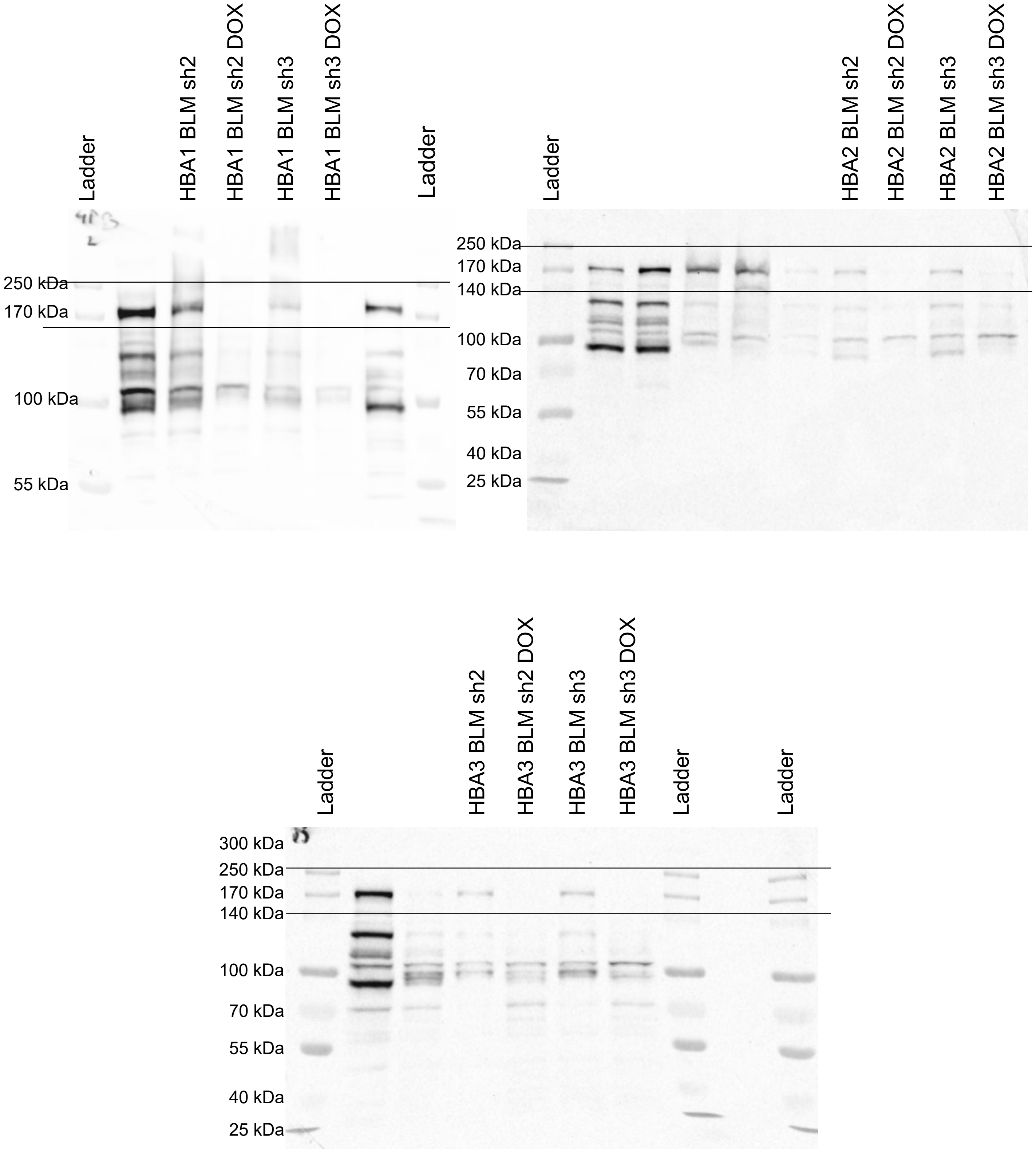


**Figure S15: Uncropped BLM western blots corresponding to Figure S8 B.**

Western blot validation of *BLM* shRNAs efficiency after 4 weeks of DOX treatment. Two different shRNAs were used in each cell line (sh2 & sh3). The Ladder used is the ProSieve® QuadColor® marker, Lonza (#00193837).

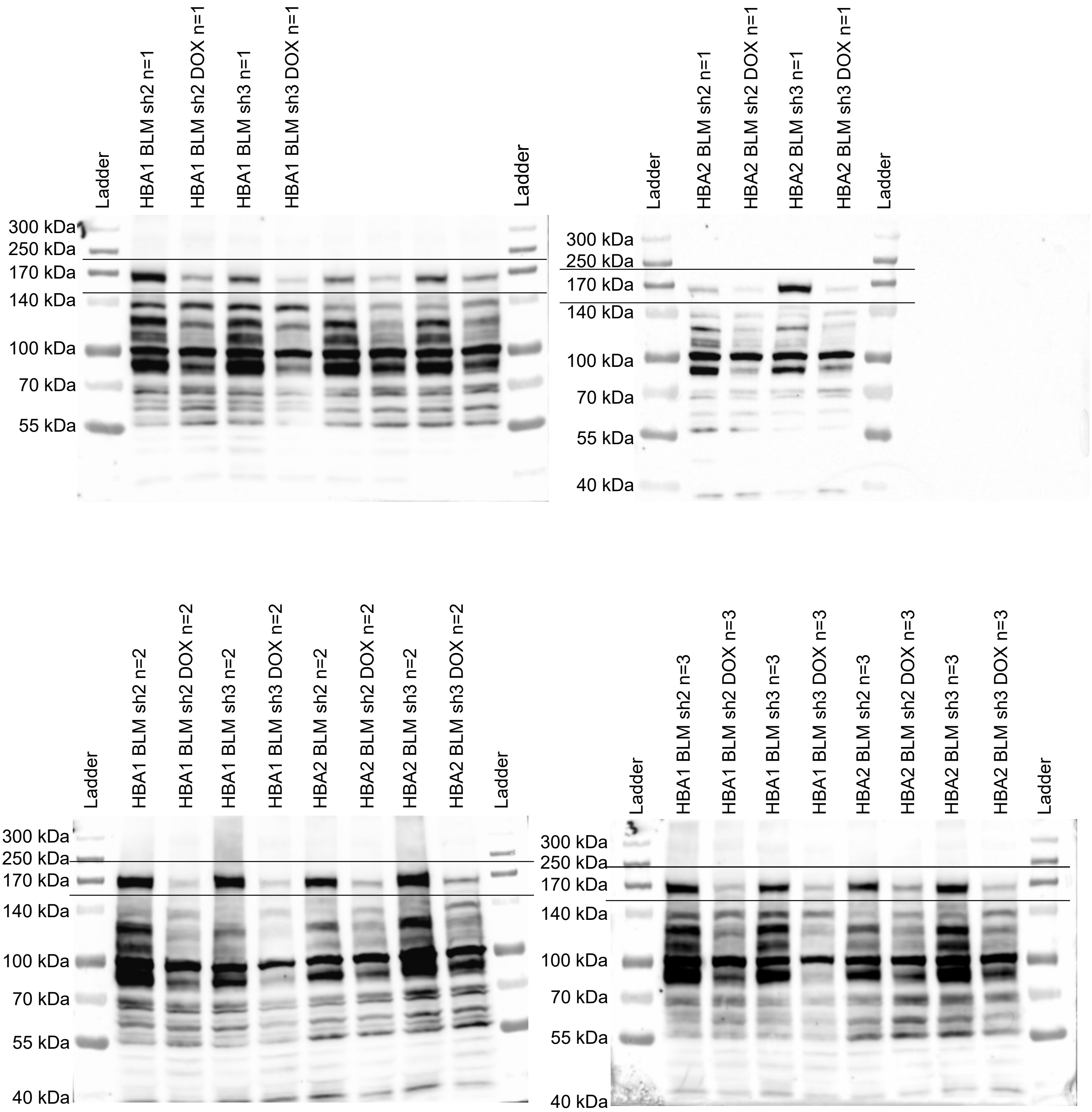


**Figure S16: Uncropped BLM western blots corresponding to Figure S12 B part 1.**

Western blot validation of *BLM* shRNAs efficiency after 5 days of DOX treatment. Two different shRNAs were used in each cell line (sh2 & sh3). The Ladder used is the ProSieve® QuadColor® marker, Lonza (#00193837).

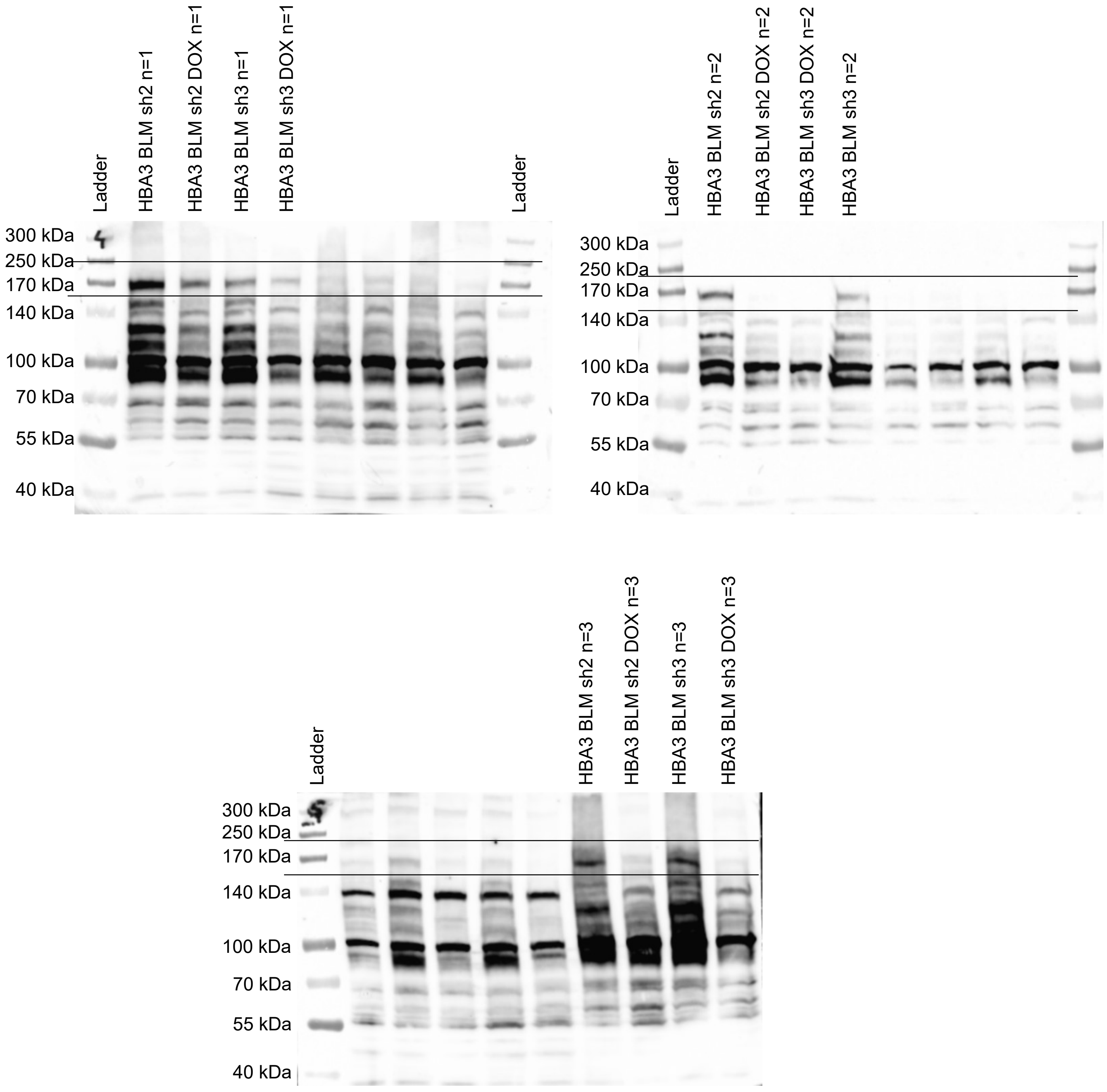


**Figure S17: Uncropped BLM western blots corresponding to Figure S12 B part 2.**

Western blot validation of *BLM* shRNAs efficiency after 5 days of DOX treatment. Two different shRNAs were used in each cell line (sh2 & sh3). In Supplementary Figure S12B, the Western blot for HBA3 (n = 2) was split into two parts, and the portion containing the HBA3 BLM sh3 DOX and HBA3 BLM sh3 lanes was inverted in order to present the HBA3 BLM sh3 condition first, followed by the HBA3 BLM sh3 DOX condition. The Ladder used is the ProSieve® QuadColor® marker, Lonza (#00193837).
